# A sensory-motor decoder that transforms neural responses in extrastriate area MT into smooth pursuit eye movements

**DOI:** 10.1101/2023.05.12.540526

**Authors:** Stuart Behling, Stephen G. Lisberger

## Abstract

Visual motion drives smooth pursuit eye movements through a sensory-motor decoder that uses multiple parallel components and neural pathways to transform the population response in extrastriate area MT into movement. We evaluated the decoder by challenging pursuit in monkeys with reduced motion reliability created by reducing coherence of motion in patches of dots. Reduced dot coherence caused deficits in both the initiation of pursuit and steady-state tracking, revealing the paradox of steady-state eye speeds that fail to accelerate to target speed in spite of persistent image motion. We recorded neural responses to reduced dot coherence in MT and found a decoder that transforms MT population responses into eye movements. During pursuit initiation, decreased dot coherence reduces MT population response amplitude without changing the preferred speed at the peak of the population response. The successful decoder reproduces the measured eye movements by multiplication of (i) the estimate of target speed from the peak of the population response with (ii) visual-motor gain based on the amplitude of the population response. During steady-state tracking, the decoder that worked for pursuit initiation failed. It predicted eye acceleration to target speed even when monkeys’ eye speeds were steady at a level well below target speed. We can account for the effect of dot coherence on steady-state eye speed if sensorymotor gain also modulates the eye velocity positive feedback that normally sustains perfect steadystate tracking. Then, poor steady-state tracking persists because of balance between deceleration caused by low positive feedback gain and acceleration driven by MT.

---

Primates use smooth pursuit eye movements to track moving objects with their eyes. The system uses visual motion to initiate and correct ongoing pursuit (Robinson 1965). It uses motor feedback to maintain eye speed and allow essentially perfect tracking (Lisberger 2010). The initiation of pursuit is an open-loop movement where the visual motion drives eye speeds before any feedback can ‘interfere’ with final eye movements (Lisberger and Westbrook 1985). The goal of pursuit is to match target speed with eye speed by the end of the open-loop interval, meaning that non-visual drivers of eye movement are needed to maintain eye speeds during perfect steady-state tracking (Morris and Lisberger 1987). Several lines of evidence implicate positive feedback of previous motor commands through the cerebellar floccular complex in sustaining eye speed through steadystate tracking (Lisberger and Fuchs 1978a, 1978b; Miles and Fuller 1975; Stone and Lisberger 1990).

Most previous studies of visual control of pursuit, including our own, have focused on using measurements of eye movements to infer how target motion is transformed into motor output and then creating models that use engineering strategies to transform target motion into eye motion (Churchland and Lisberger 2000; Orban de Xivry et al. 2013; Ringach 1995; Robinson et al. 1986). For example, our previous work revealed properties of sensory-motor processing in a control-theory model of pursuit by showing how we could account for deficits introduced by reducing the coherence of motion within a patch of dots that serves as the target (Behling and Lisberger 2020). Low dot coherence causes deficits in eye speed during both pursuit initiation and steady-state tracking. We attributed reduced eye speed during pursuit initiation to reduced strength of visual-motor transmission secondary to reduced motion reliability. However, this explanation fails during steady-state tracking when low coherence patches of dots perplexingly cause eye speed that is persistently lower than target speed even though the residual image motion should accelerate eye speed to target speed. To explain this paradox, we suggested that low motion reliability also causes reductions in the gain of eye velocity positive feedback through the floccular complex.

Our goal here is to supplant the kinematics of the target motion as the input to models of pursuit and to replace it with activity recorded in the brain area responsible for delivering signals about target speed and direction to the pursuit system. We record from the middle temporal visual area in the extrastriate cortex (MT) during the initiation and steady-state tracking of targets with reduced dot coherence. Then, we reveal models of the sensory-motor decoder that can transform the population response in MT into the eye movements we measure during both the initiation of pursuit and steady-state tracking. As a major source of the visual motion signal used for pursuit (Dürsteler and Wurtz 1988; Newsome et al. 1985; Priebe et al. 2003), area MT is the critical locus of the visual processing pathway to investigate the mechanisms that cause deficits in eye speed during pursuit of a patch of dots with low dot coherence. Neurons in area MT respond preferentially for specific directions and speeds of motion (Dubner and Zeki 1971; Lisberger and Movshon 1999; Maunsell and Van Essen 1983). They also have specific areas of the visual field where they are responsive to that motion (Desimone and Ungerleider 1986), along with complicated suppressive surrounds (Born and Tootell 1992). Based on prior studies of MT using random dots (Britten et al. 1993), it seemed likely that responses would degrade as we reduced dot coherence in our stimuli.

Our analysis recognizes that no one neuron in MT provides an unambiguous readout of the speed of target motion. Instead, the population of the neurons in MT encodes the speed of targets that drive accurate pursuit (Lisberger 2015). Because of the importance of the “population response”, the key question is whether and how the MT population response can be decoded to estimate either the physical motion of the target or the eye movements it evokes. We leverage reduced dot coherence to challenge the system and help us answer this question by searching for decoders that would transform the MT response appropriately across both target speeds and dot coherence, and also during the initiation and steady-state phase of pursuit tracking.

Our previous papers developed the concept of a sensory-motor decoder that transforms the MT population response into the initiation of pursuit based on two parallel pathways (Darlington et al. 2017; Egger and Lisberger 2022). One pathway estimates the physical image motion across the retina and the other pathway uses multiple factors including the amplitude or signal-to-noise ratio in the MT population response to control the strength (or “gain”) of visual transmission to the pursuit system. MT provides inputs to the smooth eye movement region of the frontal eye fields (FEF_SEM_) (Stanton et al. 2005) and FEF_SEM_ is responsible for control of the gain of visual transmission (Tanaka and Lisberger 2002a, 2002b). When the stimulus contains reduced dot coherence, we propose that some property of the MT population response drives FEF_SEM_ to decrease the gain of visual-motor transmission and reduce the evoked eye speed in the initiation of pursuit (Behling and Lisberger 2020). Here, we test our concept with neural data, both for pursuit initiation and to explain why low dot coherence causes steady-state eye speeds that fail to accelerate to target speed.

Here, we report the responses of a large population of MT neurons during the initiation and steadystate of pursuit for a range of dot coherences and target speed. Our analysis shows that MT speed representations are veridical in the sense that they estimate target speed correctly even for low dot coherences. The effect of dot coherence on the initiation of pursuit can be attributed to a component of the sensory-motor decoder that uses the reduced response amplitude in the MT population response to drive a lower gain of visual-motor transmission. However, the same decoder predicts that the MT population response during steady-state tracking should drive eye acceleration up to target speed, even for low dot coherences. We suggest that low dot coherence causes reduced gain in motor feedback through the cerebellum. The reduced motor feedback gain favors eye deceleration that is balanced by eye acceleration drive from MT, leading to stable steady-state eye speeds that are (no longer perplexingly) below target speed.

## Methods

We studied two male rhesus macaque monkeys between the ages of 8 and 16 years and weighing between 9 and 12 kg. Several surgeries were performed under anesthesia with IACUC approval and veterinary observation to implant (i) a head post to restrain head movement, (ii) a scleral eye coil to track eye movements, and (iii) recording chambers over area MT that could be sealed when not in use, to allow introduction of microelectrodes. Monkeys were trained to enter and sit within a primate chair and to associate a juice reward with fixating and visually pursuing targets on a CRT monitor.

### Visual Stimulus, Behavior, and Recording

CRT monitors were 23-inches diagonal (58.4 cm), were placed 30 cm from the monkey’s eyes, and had a refresh rate of 80 Hz. Targets, presented on a neutral gray background, were generated and displayed using custom lab software called “Maestro”. The targets comprised, sequentially, of black fixation dots with a diameter of 0.6 deg and patches of 72, 5-pixel-wide dots in a 4 deg diameter circular aperture; half of the dots were white and half were black. As illustrated in Figure 1B, trials started with the fixation target in the middle of the screen for 400 to 600 ms randomized over a uniform distribution. A grace period of 300 ms gave the monkey time to fixate at the target after which the eye needed to remain within 2 deg of the fixation target to allow the trial to go forward. After the fixation period expired, a patch of dots appeared with local motion of the dots at speeds of 2, 4, 8, 16, or 32 deg/s within the patch in cardinal or oblique directions for 100 ms. Then, the motion of the dots relative to the patch transitioned entirely to the global movement of the patch as a whole at the same speed. The full target motion was created as if there was a fullfield display of dots moving with a particular direction, speed, and motion coherence with an invisible aperture that is stationary over the full field display for 100 ms before moving in the same direction and speed of the dots. The use of local-then-global motion created pursuit initiation identical to that produced without the period of local motion (Osborne and Lisberger 2009), but allowed us to minimize catch up saccades during pursuit initiation and contrived for the stimulus to remain at a fixed location on the receptive field of the neuron under study for the first 100 ms of the stimulus. The size of the fixation window was increased to 4 deg during global motion. The dot patch moved for 1200 to 1400 ms randomized over a uniform distribution. At faster speeds the fixation window was occasionally increased near the end of the trial if the location eccentricity was approaching the edge of the screen. At the end of each trial, the dot patch jumped in position by 10% of target speed and stopped to help prevent additional saccades and hold for final fixation and reward.

**Figure 1:**
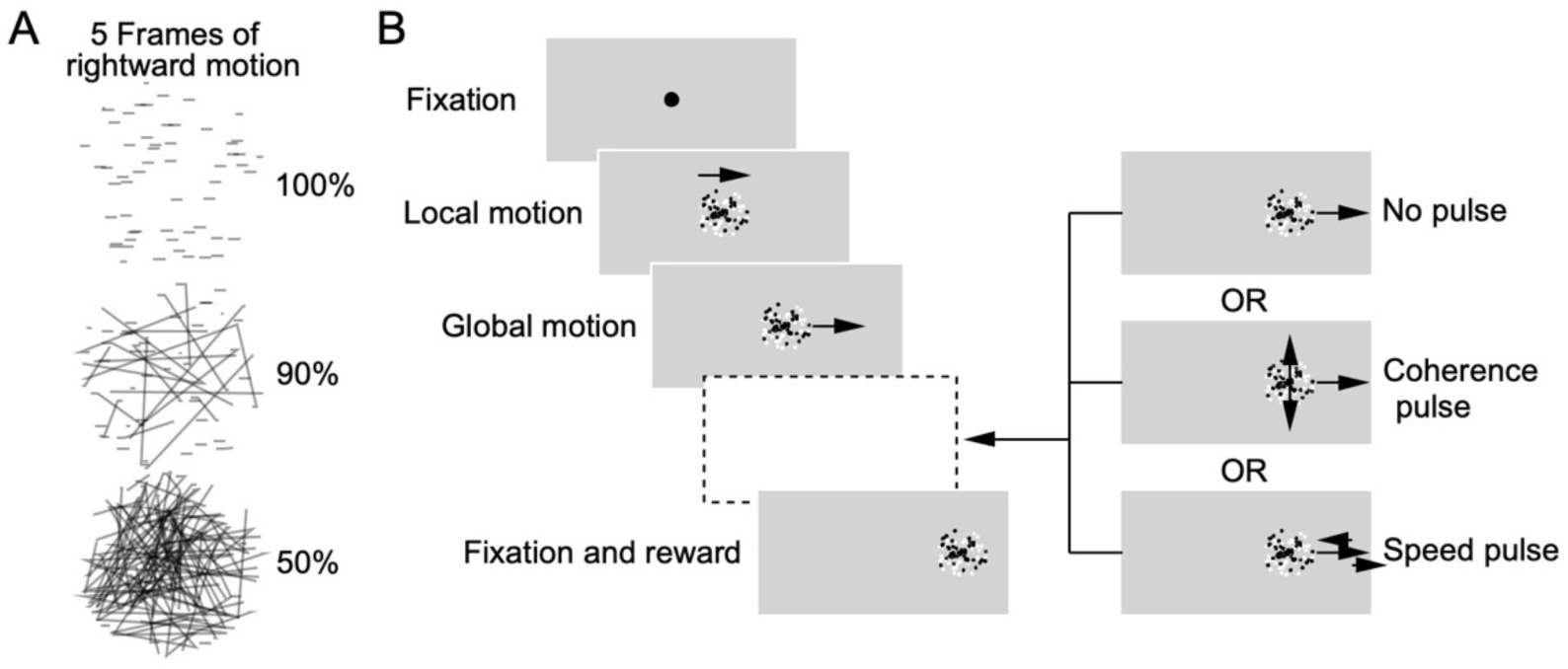
Structure of the experimental task: **A**: Examples of what motion looks like for different levels of dot coherence. For five frames of rightward motion, 100% dot coherence shows each dot in a line, 90% shows each dot with a 90% chance of staying in a line or randomly reappearing somewhere else within the aperture, and 50% only has a 50% chance of staying in a line or randomly moving to another position. **B**: The task structure with fixation, local motion of dots within the aperture, global motion of the patch of dots as a whole, continuation of motion with or without changes in speed or dot coherence, and finally fixation and juice reward.

Experiments delivered randomly-interleaved trials with different directions, speeds, and coherences within the patch of dots. While the monkeys were pursuing repeated trials of the motion of targets of different speed and dot coherence, we recorded neurons within foveal MT. We focused on isolating the neuron with the strongest responses and selected the preferred direction axis for the stimulus set from the direction tuning of that neuron, though frequently we recorded additional responsive neurons with similar tuning. We recorded eye movement responses as position and velocity signals from the eye coil. We obtained eye velocity signals with an analog circuit that differentiated frequencies below 25 Hz and filtered out higher frequencies above with gain that decreased at –20 dB/decade. Samples at 1 kHz were saved along with codes that would allow synchronization with neural data.

We used FHC single electrodes to map the recording chambers. We found and identified foveal MT based on well-documented responses to motion that were speed and direction tuned and constrained to within several degrees of the fovea of the visual field while the monkey was fixating (Desimone and Ungerleider 1986). TREC tetrodes and Plexon 16 channel S-probes then were used for the majority of the data collection and Plexon Omniplex systems were used for data recording. Every channel’s continuous wideband signal was saved for offline sorting except for several early recordings where we recorded only waveform clips of threshold crossings and spike times. Plexon offline sorter was used for sorting older files with only threshold crossing and spike times and custom sorting software and the Full Binary Pursuit spike sorter developed in our laboratory (Hall et al. 2021) was used for the majority of the recordings. Sorted spike data then was combined with behavioral trial data and synced for analysis using custom functions written in Julia.

We defined coherence as the frame-to-frame probability that a given dot continues at the defined direction and speed of the moving patch versus being moved randomly to another location within the patch (Figure 1A). It is important to note that coherence operates the same way for local motion as for global motion. Previous literature referred to the proportion of coherently moving dots as motion correlation (Britten et al. 1992), but our strategy should not be confused with direction noise where proportions of dots move in different directions but at the same speed. In our stimulus, coherence manipulates stimulus strength through the distribution of motion energy around the global speed at the center of the distribution (Britten et al. 1993). For example, at 0% coherence, the location of each dot is randomly changed each frame; the random individual motion events for each dot do not create net bias that competes with the global motion. At values of coherence lower than 100%, the individual, disparate motion events collectively degrade and redistribute the motion energy around a small central peak created by the movement of the aperture. Degrading motion without removing the information about target speed is precisely our goal by changing dot coherence.

### Simulated Model Population

We used the sum of a series of Gaussian function curves with peaks at different times after the onset of target motion to model our neurons’ activity. The individual Gaussians were modeled as:

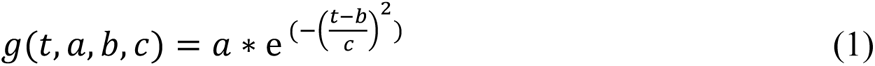

where *t* represents time in ms and *a*, *b*, and *c* represent the amplitude, the center, and the width of each Gaussian. The full “sum-of-Gaussians” model *M* was:

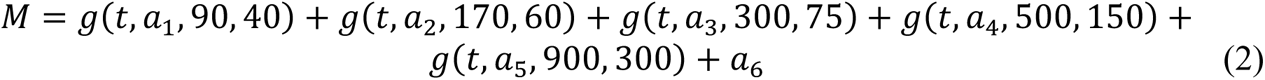

We used an iterative procedure to end up with sets of parameters that allowed us to fit well the responses of each neuron as a function of stimulus speed and dot coherence. First, we chose initial parameters for each neuron based on the average of its time varying firing rates during pursuit across stimulus speeds but fitting separately for pursuit in the preferred and null directions and separately for 100%, 30%, and 10% dot coherences. Next, we fitted Equation (2) to the responses at each dot coherence and each target speed, with the goal of determining how the amplitudes in Equation (2) need to change as coherence decreases and as the difference between preferred speed and target speed increases. The fits to individual speeds and coherences yielded a good model output for each neuron and allowed us to predict the response at any speed or coherence by linearly interpolating between the values of a_1-5_ for any stimulus speed or dot coherence. This model has many parameters (n=36), but it reproduces the time varying firing rates of the MT population well and therefore provides an excellent database as an input to multiple models of the sensory-motor decoder for pursuit. In general, the values of a_1-5_ behaved sensibly as functions of target speed and dot coherence.

To create a homogeneous model population with a number of model neurons much larger than our sample, we drew randomly from a uniform distribution of preferred speeds to select the preferred speed of the model neuron. We selected a response profile by drawing randomly from the 100 neuron models where the speed tuning curve had a peak that was at neither the top nor the bottom of the range of speeds we tested. We then created model responses based on the parameters of the fits to the responses of the recorded neuron but with the randomly chosen preferred speed. For each model neuron, we simply slid the linear interpolation for the three coherences slower or faster to correct the difference between the preferred speed of the recorded neuron and the randomly chosen preferred speed of the model neuron. To obtain the responses of the model neuron at the target speeds we used to decode the resulting model population response, we adjusted the amplitudes of the sum-of-Gaussian model fits according to the new interpolated values. This last step would not have worked well for neurons that didn’t have a peak in the range of speeds we tested, thus our exclusion criteria.

### Decoding and analysis

The decoding equations we used were based on opponent vector averaging:

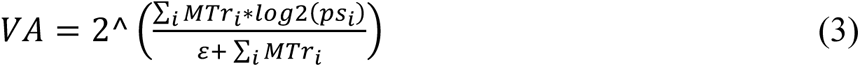

where *MTr_i_* is the normalized opponent response of the *i*^th^ MT neuron, real or simulated, and *ps_i_* is the preferred speed of the *i*^th^ neuron. The anti-noise term *ε* prevents the decoded values from exploding when firing rate is low, and is treated as a constant based on (Priebe and Lisberger 2004).

Decoding by gain modulation of the vector average of MT was calculated as:

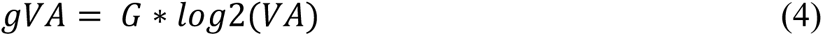

For the gain modulated vector averaging decoder, the gain was calculated to represent the overall amplitude of the real or simulated MT population response as:

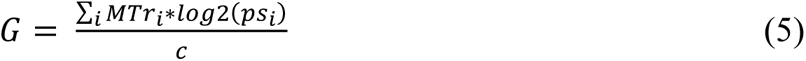

where *c* is a term based on the size of the population and the average value for ideal decoding of each speed in the stimulus set.

Our statistical testing used Welch’s t-test for unequal variance. Even though normality and equal variance were frequently violated in our data, this test covers type 1 errors (rejecting a true null hypothesis) better than some non-parametric alternatives (Zimmerman 1998). We also used kernel density estimations for Mann-Whitney U tests for building and for comparison of the simulated population responses with the data responses when the population sizes were dramatically different.

## Results

Our database included 203 well-isolated and responsive MT neurons with receptive fields on or very close to the fovea, 92 from monkey Ar and 111 from monkey Di. We recorded their responses to moving targets during pursuit initiation and steady-state tracking. Our primary stimulus was a patch of dots with different values of motion coherence with the motion selected from a range of directions and speeds. For studying the initiation of pursuit, the most important feature of our stimuli was that the dots moved locally within a stationary patch for the first 100 ms of motion, keeping the patch itself stable on the receptive field of the neurons under study and preventing catch up saccades while the eyes began to pursue.

Our goal is to understand the sensory basis for previous behavioral observations on the effect of dot coherence on smooth pursuit eye movements, and to reveal how sensory representations are transformed in a sensory-motor decoder to reproduce the measured eye movements. During the initiation of pursuit for 100% coherent dot motion, eye speeds rise to match target speed and are maintained for stable tracking during the steady state of pursuit. As dot coherence decreases, the initial rise in eye speeds slows and the maintained eye speeds also fall short of target speed with a slight tendency to decay towards zero (Figure 2A, B). Our question – why eye speeds decline as we reduce dot coherence – might be answered by changes in the peak or amplitude of speed tuning curves in MT or may implicate specific downstream decoding mechanisms.

**Figure 2:**
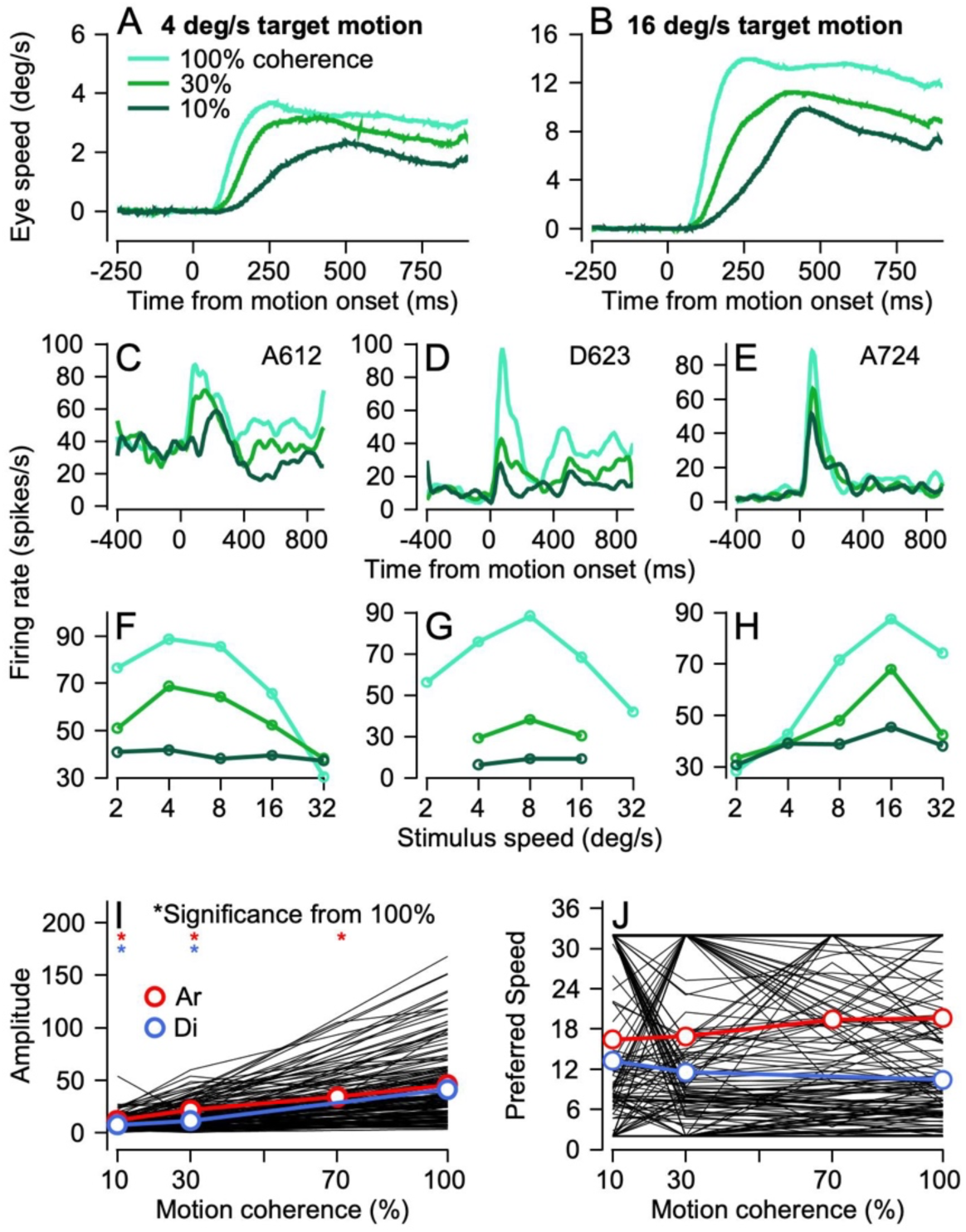
Effect of dot coherence on pursuit eye movements and responses of individual MT neurons. **A, B**: Average eye speeds for target speeds at 8 (**A**) and 16 (**B**) deg/s and 100%, 30%, and 10% dot coherences. Time 0 is aligned to onset of local motion. **C-E**: Average firing rates in spikes per second from 3 example MT neurons for stimulus motion in each’s preferred speed and direction for 100%, 30%, and 10% dot coherence. **F-H**: Speed tuning curves for stimulus motion in the preferred direction as a function of dot coherence for the 3 example neurons. In **A-H**, the colors of the traces shift from cyan to green to dark green for 100%, 30%, and 10% dot coherences. **I**: Amplitude of the tuning curves as a function of dot coherence. **J**: The preferred speed from the tuning curve as a function of dot coherence. In **I** and **J**, each black line shows data for a different single neuron and the bold red and royal blue lines with symbols show the averages across all MT neurons recorded in monkeys Ar and Di. Asterisks indicate conditions where the averages differed significantly from those at 100% dot coherence.

### Effect of dot coherence on the speed tuning curves of individual MT neurons during pursuit initiation

MT neurons respond during the initiation of pursuit with a brief transient pulse of firing rate that starts 40-60 ms after the onset of stimulus motion, followed by modest elevation of firing rate for the remainder of the trial (Figure 2C–E). The amplitude of the transient pulse decreases as a function of the coherence of the motion in the dot patch. To compute speed tuning curves during the initiation of pursuit, we measured the mean firing rate for each stimulus speed and dot coherence in the interval from 60 to 120 ms after the onset of motion. Plotting the responses for each dot coherence as a function of stimulus speed revealed clear speed-tuning curves and allowed us to identify a preferred speed for each neuron.

For the 3 example neurons in Figure 2, the relative preference for different target speeds was invariant with dot coherence. The 3 speed-tuning curves showed clear peaks at 4, 8, and 16 deg/s for 100% coherence dot patches (Figure 2F–H). When the motion coherence in the dot decreased, the amplitude of the speed tuning curve decreased but the peak of the curve, representing the preferred speed, did not shift for these individual neurons or for the majority of our sample.

Analysis of the full population of neurons from both monkeys (black lines in Figure 2I, J) reveals a consistent effect of motion coherence on the amplitude of the tuning curve, estimated as the difference between the largest and smallest responses across the speeds we presented. We used the difference as a metric of tuning amplitude to avoid the challenges of fitting tuning curves to data that were very weakly tuned at low dot coherences. Most neurons (black lines) and the averages across neurons for each monkey separately (colored lines) showed increases in tuning amplitude as a function of dot coherence. Welch’s t-tests revealed significant (p<0.05) differences between the preferred speed amplitudes at 100% coherence compared to both 30% and 10% dot coherence for both monkeys and between 100% and 70% coherence for monkey Ar.

The effect of dot coherence on preferred speed, defined as the stimulus speed that elicited the largest response, was more variable (Figure 2J). Yet, there was no consistent trend of changing preferred speed as dot coherence changed; the population values of preferred speed (Figure 2J, colored symbols and lines) do not show statistically significant effects of dot coherence. Many individual neurons (Figure 2J, black lines) showed no effect of dot coherence on preferred speed, but some individual neurons seemed to change preferred speed dramatically sometimes moving from preferring 32 deg/s to 2 deg/s and back to 32 deg/s as coherence changed from 100% to 30% to 10%. Since both those values are at the limit of target speeds we presented, we suspect that the observed chaos results from the challenges of estimating preferred speed for low neural response amplitudes.

We looked more closely at the data for individual neurons and verified that most of the apparent effects of dot coherence on preferred speed were the result of low amplitude tuning curves that made it difficult to obtain a valid estimate of preferred speed. We utilized a metric that tested statistically for each tuning curve whether the responses were different between the target speeds that had the lowest versus highest responses on the speed tuning curve. This metric, which we will call the “modulation statistic” gives a sense of whether the responses actually are tuned or, instead, should be considered to be a flat line essentially at noise level. When we pull out the neurons that had significant modulation statistics (p<0.05) over the spread of stimulus speeds we tested and across dot coherences, we see that the preferred speeds for these neurons change very little as a function of dot coherence (Figure 3A–C). There are a few exceptions with significant modulation statistics across all 3 dot coherences that did show increased or decreased preferred speeds as dot coherence decreased, for example the tuning curves illustrated in Figure 3G–J. Among the neurons with a shift in preferred speed, some increased and some decreased preferred speed with no bias in decreased preferred speed to match the behavior; the majority of preferred speeds did not shift by more than a few deg/s.

**Figure 3:**
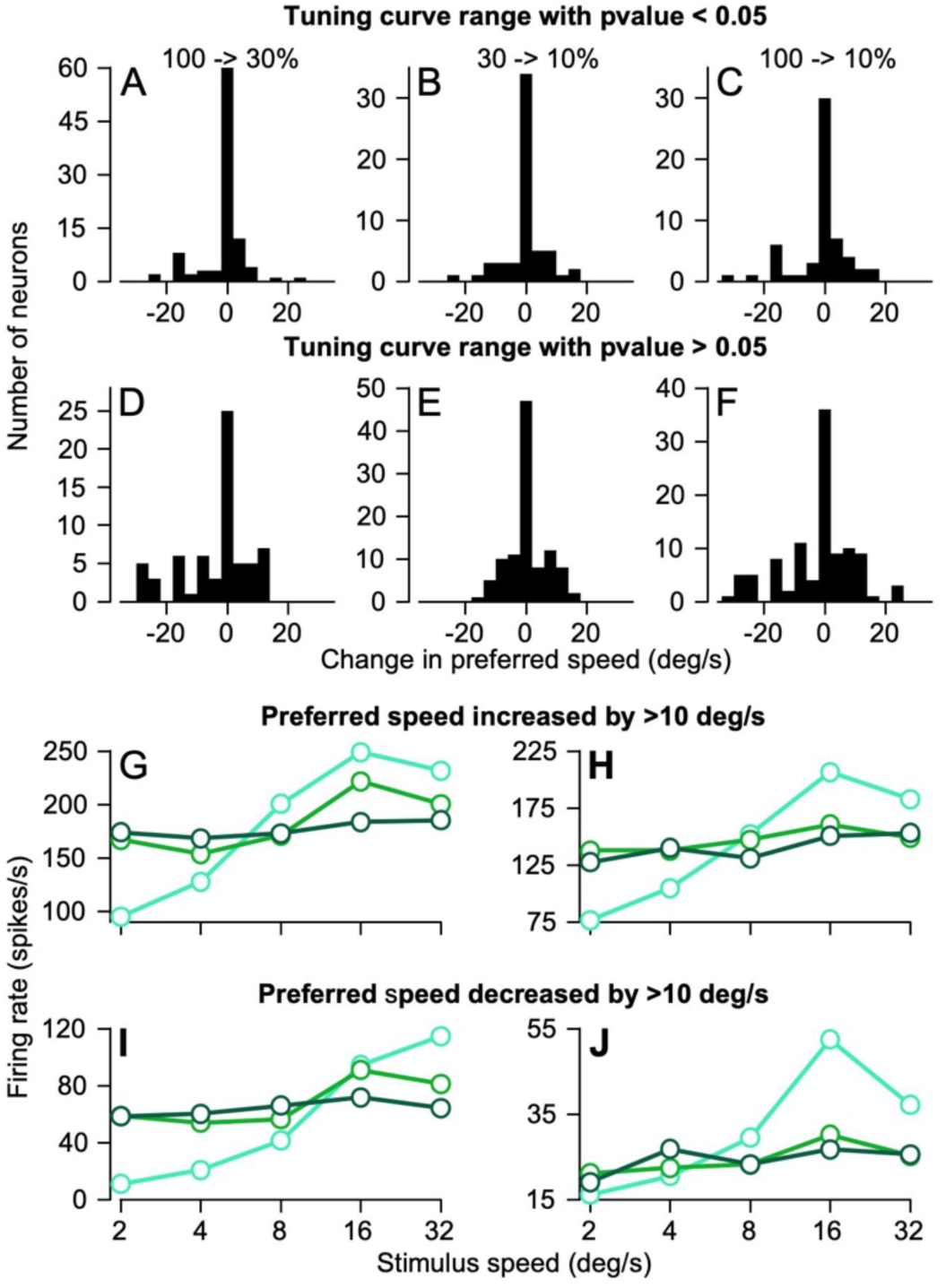
Assessment of statistical validity of apparent changes in preferred speed as a function of dot coherence. **A, B, C**: Histograms showing how much the preferred speed of each neuron changes when the tuning curve range of the lower dot coherence was statistically significant under a Welch’s T-Test with a p-value less than 0.05. Comparisons are between speed tuning curves at 100% versus 30% dot coherences (**A**) between 30% and 10% (**B**) and between 100% and 10% (**C**). **D, E, F**: The same analysis as in **A-C** but for neurons where the tuning curve range was not found to be statistically significant. **G, H**: Example speed tuning curves for 2 neurons where the preferred speed increased by at least 10 deg/s from 100% to 10% dot coherence. **I, J**: Example speed tuning curves for 3 neurons where the preferred speed decreased by at least 10 deg/s. In **G-J**, cyan, light green, and dark green show curves for dot coherences of 100%, 30%, and 10%.

Neurons whose speed tuning curves did not show significant values of modulation statistic (Figure 3D–F) had more and larger shifts in preferred speed as dot coherence decreased but again showed balanced increases and decreases in speed. We conclude that decreased dot coherence does not alter the preferred speed of MT neurons, but that some speed tuning curves become too noisy to assess preferred speed because response amplitude decreased to the point of the curve becoming a flat line. Because our full sample of neurons represents how dot coherence affects the MT population response during our pursuit task, we retained all recorded neurons in the population for all further analyses.

### Effect of dot coherence and stimulus speed on the population response in MT during pursuit initiation

The pursuit system decodes the response of the population of neurons in MT to identify the speed of target motion and determine the appropriate change in eye velocity at the initiation of pursuit. Thus, it is relevant to assess the effect of target speed and dot coherence on the MT population response through plots like those in Figure 4. Here, each neuron appears as a separate symbol and its response for a give set of parameters is plotted as a function of the neuron’s preferred speed for 100% coherence dots. To assess the population response during pursuit initiation, we measured firing rate in the interval from 60 to 120 ms after the onset of target motion. Firing rates for each neuron are normalized relative to the peak firing for target motion in the preferred direction and the opponent response is computed by subtracting identically-normalized responses to target motion in the null direction.

**Figure 4:**
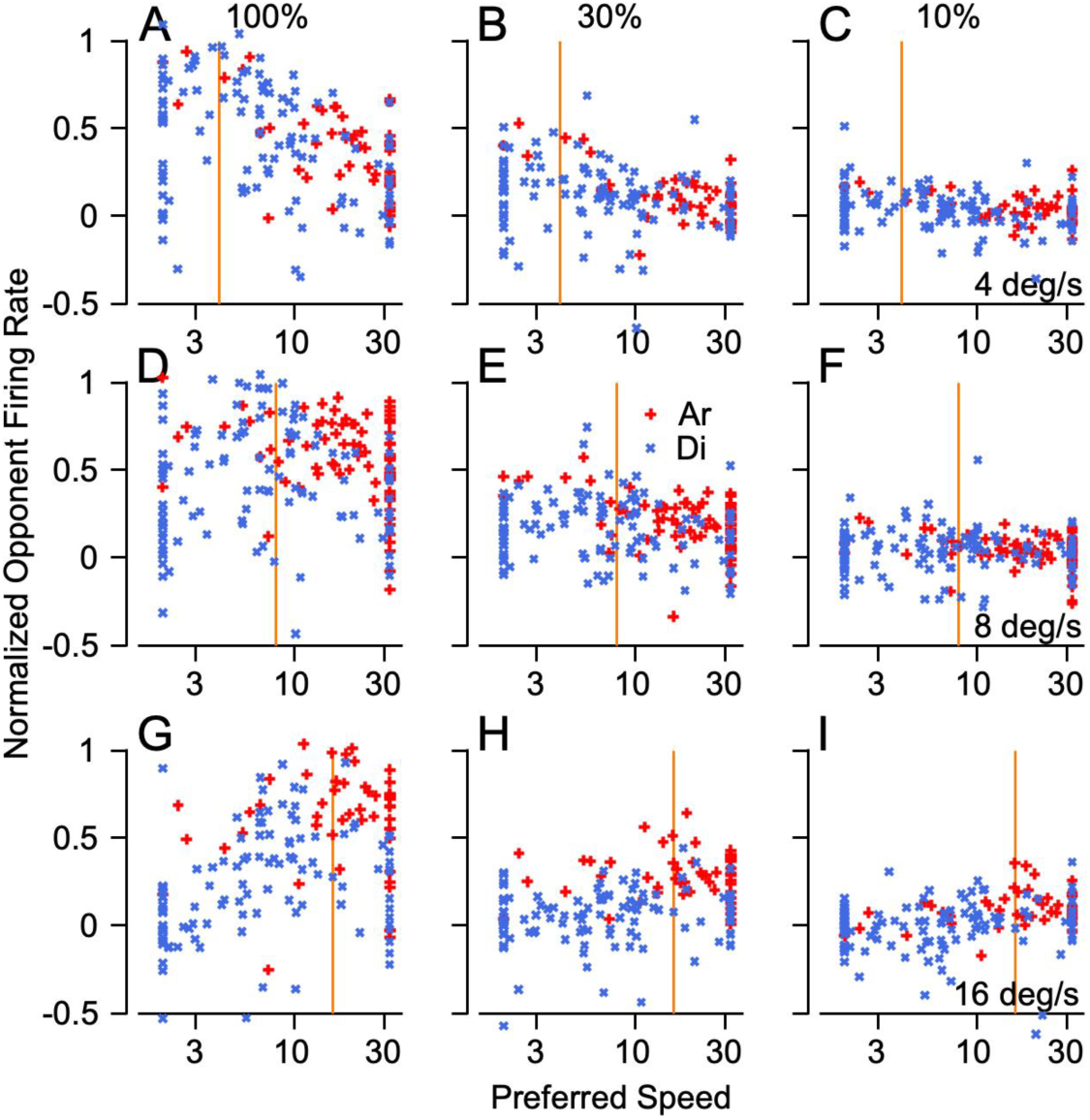
MT population responses during pursuit initiation as a function of target speed and dot coherence. In each graph, the different symbols show the responses of different MT neurons and plot their normalized opponent firing rates as a function of their preferred speed. Royal blue x’s and red crosses showing data from monkeys Di and Ar. The vertical orange lines represent stimulus speed. **A-C**: Responses for stimuli moving at 4 degrees per second at 100% (**A**), 30% (**B**), and 10% dot coherence (**C**). **D-F**: Responses to stimuli moving at 8 deg/s at 100% (**D**), 30% (**E**), and 10% dot coherence (**F**). **G-I**: Responses to stimuli moving at 16 deg/s at 100% (**G**), 30% (**H**), and 10% dot coherence (**I**).

For 100% dot coherence, the population responses (Figure 4A, D, G) show clear peaks in neurons with preferred speeds near target speed (vertical orange lines), as expected given the tuned responses of MT neurons. The same general tendency appears as dot coherence is reduced but becomes less pronounced because most neurons’ response amplitude declined as dot coherence was reduced to 30% (Figure 4B, E, H) and 10% (Figure 4C, F, I). Even for the low values of dot coherence, the responses of neurons with preferred speeds closer to the image motion from target speed were at least slightly larger than the responses of neurons with preferred speeds that were much faster or slower. Neurons from both monkeys show similar responses even though we sampled more neurons that preferred faster speeds from monkey Ar (red symbols) and more neurons that preferred slower speeds from monkey Di (blue symbols). The number of points is not identical across graphs because not every neuron from monkey Ar was recorded during every combination of dot coherence and stimulus speed because of trial and experiment constraints at the time.

### Strategies for decoding the MT population response to identify stimulus speed versus eye speed in pursuit initiation

Before asking how the responses of the MT population can be transformed to generate the initiation of pursuit, it is important to recall that eye speed during pursuit initiation has a non-linear relationship to image speed and dot coherence. We characterized that relationship by measuring eye speed 200 ms after the onset of target motion and plotting it as a function of stimulus speed (Figure 5A). In general, eye speed is less than stimulus speed. We then asked how to decode the MT population responses to reproduce stimulus speed, eye speed, or both.

**Figure 5:**
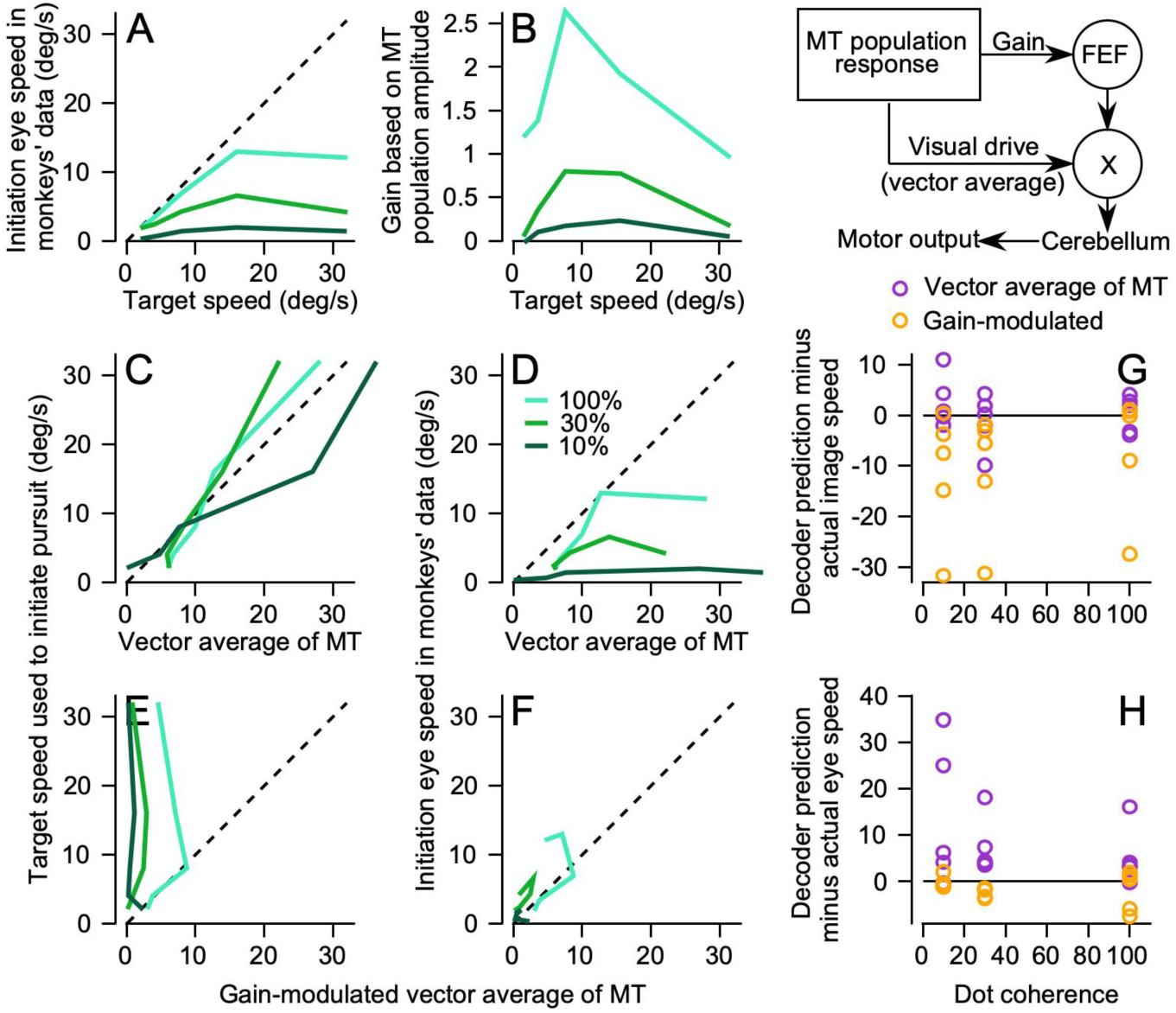
Decoding the MT population response to reproduce eye speed during pursuit initiation. The diagram in the upper right shows our conceptual framework for the flow of sensory information where the command for pursuit initiation results from multiplication of visual drive from estimating target speed by the gain of visual-motor transmission through FEF. **A**: Initiation eye speed in monkeys’ data as a function of the target speed for 3 dot coherences (100%, 30%, and 10%). **B**: The gain of visual motor transformation based on MT population response amplitude as a function of target speed. **C, E**: The target speed used to initiate pursuit as a function of decoding by a vector average of the MT population response (**C**) and as a function of decoding by gain-modulated vector average of MT (**E**). **D, F**: Similar to **C** and **E** except that the initiation eye speed in monkeys’ data is plotted as a function of vector average of MT (**D**) and gain-modulated vector average of MT (**F**). In **A-F**, cyan, green, and dark green lines show data and decodings for 100%, 30%, and 10% dot coherences. **G**: The difference between image speed and predictions of the two decoders. **H**: The difference between eye speed and the predictions of the two decoders. In **G** and **H**, purple and orange symbols show results for decoding based on an opponent vector average of MT versus a gain-modulated vector average. Multiple symbols at each dot coherence report data from multiple target speeds.

Traditional decoding strategies propose an “opponent vector average” of MT responses (see ***Methods***) to estimate the preferred speed at the center-of-mass of the MT population response (Churchland and Lisberger 2001a). If the real decoder follows the traditional strategy, then we would expect the opponent vector average to predict the observed non-linear relationship of eye speed to stimulus speed and dot coherence. Instead, the opponent vector-averaging decoder predicts an output that is very close to actual stimulus speed regardless of target speed or dot coherence (Figure 5C). As a consequence, the decoder consistently overestimates eye speed (Figure 5D). We conclude, in agreement with previous studies that degraded stimulus contrast (Krekelberg et al. 2006; Pack et al. 2005), that deficits in eye speed due to low dot coherence probably do not occur simply because area MT underestimates target speed.

A more complicated sensory-motor decoder based on previous work from our lab (Egger and Lisberger 2022) predicts eye speed well (and stimulus speed poorly) across target speeds and dot coherences. The elaborated decoder is based on data about a node in the pursuit circuit downstream from MT, namely the smooth eye movement region of the frontal eye field (FEF_SEM_). FEF_SEM_ receives population activity from MT and appears to determine, encode, and control the strength (or gain) of visual-motor transmission for pursuit. Previous papers from our laboratory (Darlington et al. 2017, 2018; Egger and Lisberger 2022) demonstrate multiple ways that models reproduce pursuit eye movement behavior if they are based on a decoder with multiple pathways based on the sketch in the upper right panel of Figure 5. One pathway uses opponent vector averaging of the MT population response to identify stimulus speed. The other pathway uses the amplitude of the MT population response to set the gain of visual-motor transmission. To test the “gain modulated” vector averaging model, we calculated gain as the summed opponent firing rates in the MT population weighted by preferred speed and divided by a constant based on population size. The resulting gain has a non-linear relationship with target speed and decreases with lowered dot coherences (Figure 5B).

When its inputs come from our MT recordings, the “gain-modulated vector average decoder” underestimates stimulus speed (Figure 5E) but predicts well the eye speed in the initiation of pursuit across values of stimulus speed and dot coherence (Figure 5F). Figure 5G and H summarize the differential performance of the vector averaging decoder versus the gain-modulated vector averaging decoder by plotting the differences between each decoder’s output and image speed (G) or eye speed (H). The vector average decoder (purple symbols) predicts target speed fairly well (G) but fails to predict eye speed (H). The gain-modulated vector average decoder (orange symbols) predicts eye speed well (H) and fails to predict target speed (G).

Decoding the recorded MT population response supports our conclusion that gain-modulation of a vector averaging decoder is necessary to account for the effects of stimulus speed and dot coherence on eye speed in the initiation of pursuit. However, there are some caveats. First, our recorded population is small compared to the actual number of neurons in area MT. Second, we used target speeds from 2 to 32 deg/s and so our population is limited to preferred speeds between 2 and 32 deg/s whereas MT contains many neurons that respond, and have preferred speeds, outside of the range we used. Below, we address these issues by generating a larger simulated population that evenly tiles a larger range of preferred speeds.

### Relationship between the speed tuning of MT neurons during pursuit initiation and their responses during steady-state tracking

Our first approach to understanding the output of MT during steady-state tracking asked whether a simple speed-tuning-curve model based on the responses of MT neurons during the initiation of pursuit also can predict their responses during steady-state tracking. If the relationship between firing rate and image speed persists during steady-state tracking, then we should see population peaks around the image speed we measured during steady-state tracking at each dot coherence, as in Figure 4. To create MT population responses during steady-state tracking, we computed a normalized opponent response as a function of time for each neuron by subtracting the null direction responses from the preferred direction responses, all normalized to the peak firing for motion of 100% coherence dots in the preferred direction. Then we fitted each neuron’s tuning curve during pursuit initiation with a Gaussian curve at each dot coherence. We created a simulated population response by resampling from the pool of speed tuning curve fits with replacement 2058 times. We assigned a preferred speed to each of the 2058 model neurons by drawing from a uniform distribution of 1 to 64 deg/s in intervals of 0.125 in log base 2. We shifted each tuning curve to align to the selected preferred speed while keeping the height and width of the tuning curves constant, the latter in log_2_ coordinates. We then used image speed to predict the normalized firing rate from each model neuron in the simulated MT population for the initiation of pursuit or during steady-state tracking.

The data and the simulated population overlap reasonably well during pursuit initiation for patches of dots with both high (Figure 6A, C) and low coherence (Figure 6B, D). However, the same is not true during steady-state tracking (Figure 6E–H). The most egregious failure of the speed tuning curve during steady-state tracking is for motion of 100% coherence dots at 16 deg/s (Figure 6G), where the actual responses during steady-state tracking (symbols with black outlines) lie well below the predictions of the speed-tuning curves (low transparency blue symbols). Other speeds and dot coherences (Figure 6E, F, H) showed smaller but still noticeable disagreements between the actual responses and the predictions from the speed tuning curves, especially for the larger values of preferred speed.

**Figure 6:**
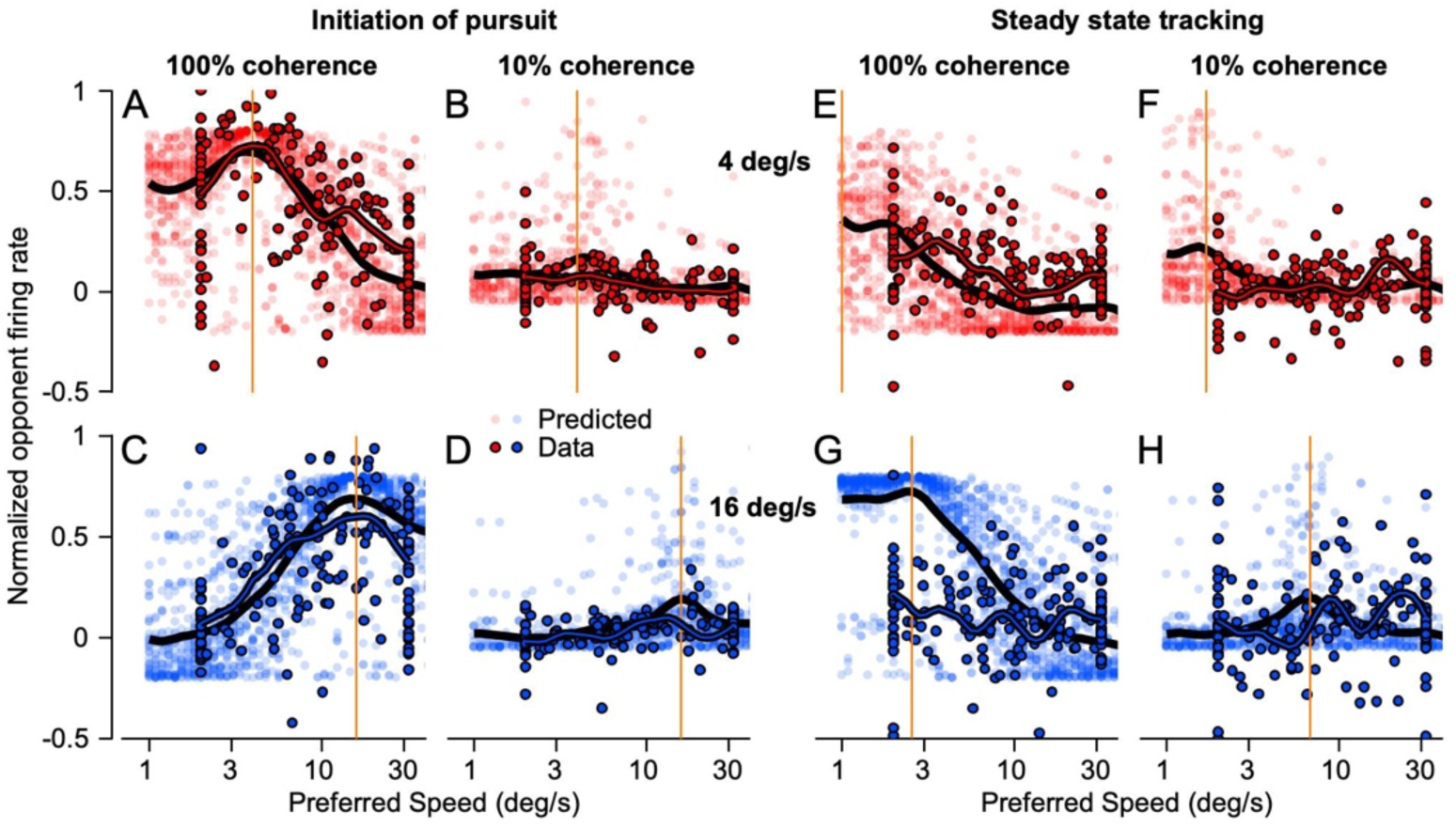
Speed tuning curves for pursuit initiation fail to predict MT responses during steadystate tracking. In these 8 population response graphs, each bold symbol plots data for an MT neuron and shows normalized opponent firing rate as a function of preferred speed. The faded symbols plot the responses of 2058 model neurons used to create a sanitized MT population response based on the speed tuning curves during pursuit initiation. **A-D**: Initiation of pursuit. **E-H**: Steady-state tracking. Red versus blue symbols are for target speeds at 4 deg/s versus 16 deg/s in the top or bottom row of graphs. **A, C, E, G**: 100% dot coherence. **B, D, F, and H**: 10% dot coherence. In each graph, the thick black line is the population average of the model’s responses and the thick red or blue line outlined in black is the population average of the data responses. The vertical orange lines indicate the image motion that drove each population response.

We conclude that the response of MT neurons become modified by the more complicated image motion of the target and background that occurs as the eye moves across the stationary scene during steady-state tracking. For any meaningful analysis of steady-state tracking, a simulated population needs to recapitulate observed MT activity much more faithfully.

### A “sum-of-Gaussians” model that reproduces MT neuron responses across the full pursuit response

Our next strategy to generate a simulated population response was to find a model that could reproduce the firing rates of MT neurons for the whole trial. Here, our goal was not to create a mechanistic model of MT responses, largely because we did not have access to structure of each neuron’s classical receptive field or suppressive surround. The responses we measured during steady-state tracking probably reflect the image motion of the target across the retina convolved with the non-classical effects of oppositely-directed image motion of the background across the retina during sustained eye motion. Instead, our goal was to create a model of a realistic MT population response by fitting the responses of MT neurons as a function of time during both the initiation and steady-state of pursuit across target speeds and dot coherences. We then could create a larger, evenly distributed model population response for the entire duration of smooth pursuits using the strategy outlined in the previous section, but now drawing examples from the more accurate fits.

For each neuron in our sample, we fitted the responses across target speeds and dot coherences with a “sum-of-Gaussians” model based on the sum of a series of five Gaussian functions (Figure 7A) with the center and width of each Gaussian fixed at values selected from trends within the average population. The free parameters for each neuron were the amplitudes of the five Gaussians, the baseline firing rate, and functions that related each Gaussian’s amplitude to target speed and dot coherence (see ***Methods***). We fit the amplitudes to the time-varying firing rates across target speed and dot coherence and normalized the responses relative to the that for the preferred speed during pursuit initiation at 100% dot coherence. Our fitting procedure worked well, as evidenced by the comparison of actual firing rates and model fits for an example neuron (Figure 7B). Because our procedure for creating a model MT population response required neurons that were explicitly tuned, we excluded neurons with preferred speeds at the highest and lowest speeds presented, leaving 100 neurons. We performed the same procedure for responses to target motion in the null direction so we could calculate opponent responses as we did with the data.

**Figure 7:**
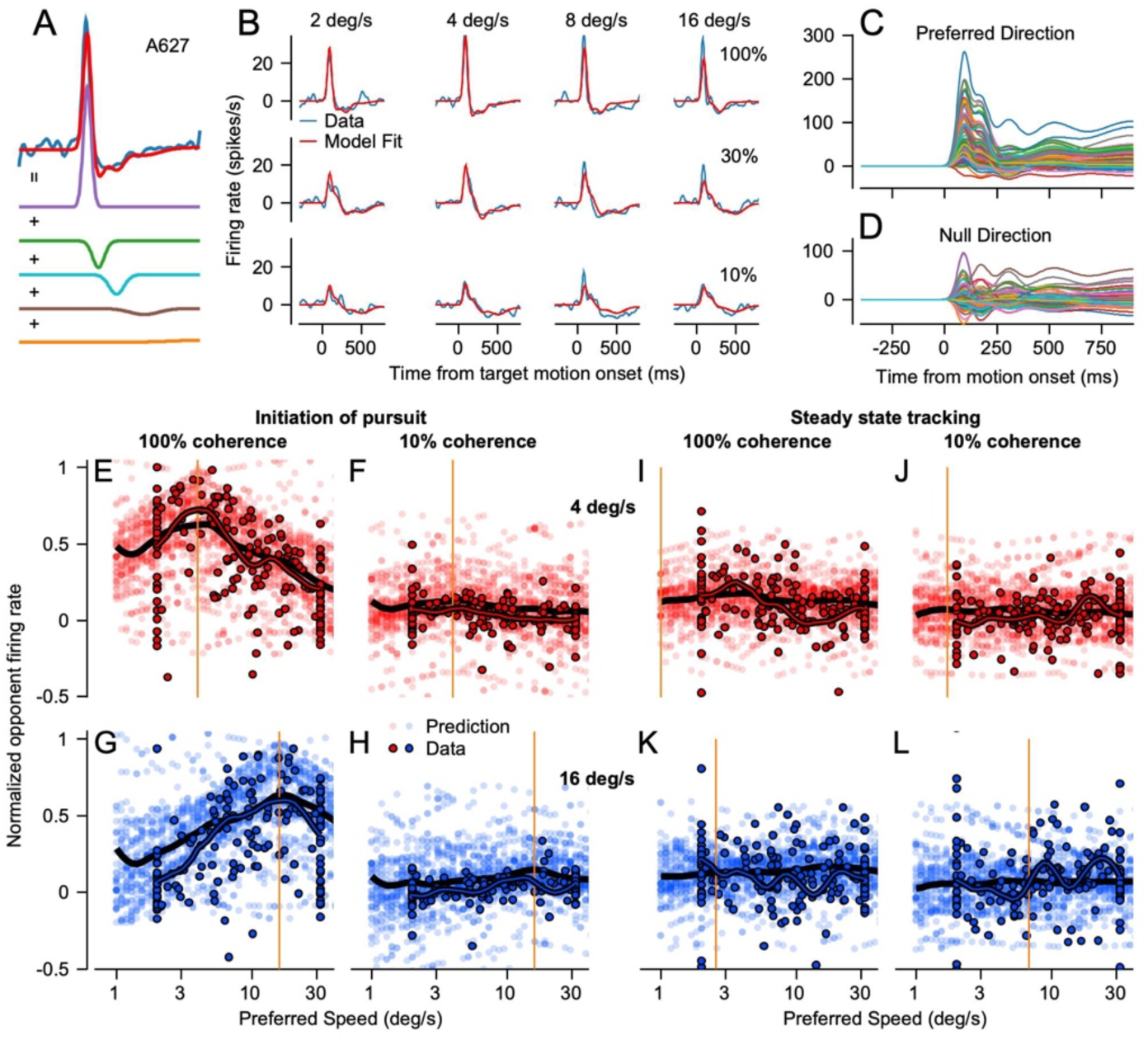
A sum-of-Gaussians model to reproduce the responses of MT neurons during pursuit initiation and steady-state tracking. **A**: Example showing how 5 Gaussians with different widths summed to reproduce a single response of an MT neuron. Red and blue traces superimposed at the top of the panel show the prediction of the model and the smoothed data. **B**: Example model fits in red superimposed over the data in blue. The columns are for target speeds 2, 4, 8, and 16 degrees per second and the rows are for dot coherences of 100%, 30%, and 10%. **C, D**: Superimposed time-varying model fits for all 100 neurons in the preferred (**C**) and null direction (**D**). **E-L**: Population responses plotted in the same fashion as in Figure 6 for predicted activity based on the sum-of-Gaussians fits to the data.

Once we had the 100 templates (Figure 7C, D) we created a model population response with a variation of the same procedure used before. We drew 2058 sets of templates at random from our 100 neuron fits and randomly assigned each set with a preferred speed from the range of 1 to 64 deg/s in intervals of 0.125 in log base 2. We then shifted the speed tuning of each set of templates so that it matched the randomly assigned preferred speed. After shifting, we had to interpolate and extrapolate to generate model responses to the specific target speeds in our stimulus set. For target speeds that fell within the five speeds we tested, we interpolated between the parameters of the model. For target speeds that fell outside the five speeds we presented, we computed the average change in each amplitude parameter across the speeds we did have fits for and used that estimate to extrapolate in log_2_ coordinates to create responses to the target speed.

The model MT population responses based on the sum-of-Gaussian fits to the data agreed well with measurements from the actual data (Figure 7E–L). Here, we averaged each model neuron’s response in intervals from 60-120 ms after onset of target motion for pursuit initiation and from 660-720 ms after onset of target motion for steady-state tracking and plotted the measurements as a function of the preferred speed of the model neuron. The measurements from simulated MT neurons (low transparency symbols) agree well with the actual data from recorded neurons (symbols with black outlines) during both pursuit initiation and steady-state tracking for 4 and 16 deg/s targets at dot coherences of both 100% and 10%. The population averages, portrayed as thick black lines for the simulated neurons and outlined in black for the recorded neurons, highlight the similarities between the simulated and recorded population responses as well as the deviations. Large, simulated MT population responses like the ones in Figure 7E–L recapitulate the recorded MT activity throughout pursuit and have greater generalizability than the actual data. Thus, our model MT population will provide inputs for future biologically-realistic models of the sensorymotor decoder.

### The sum-of-Gaussians model population behaves as well as the actual data during pursuit initiation

We validated the accuracy of the MT model by demonstrating that decoding based on the model MT population responses agreed well with decoding based on the actual data. We made it possible to predict eye speed across target speeds we were not able to present and derive smoother curves by fitting the eye speed data (Figure 8A, symbols) with a linear plateau function and extrapolating to target speeds from 1 to 48 deg/s. Opponent vector averaging of simulated MT responses predicted target speed fairly well across dot coherence, though perhaps less well for 100% coherence dots (Figure 8D). In agreement with the decoding from the recorded MT population responses, the decoder based on gain-modulation of opponent vector averages recapitulated eye speed (Figure 8E).

**Figure 8:**
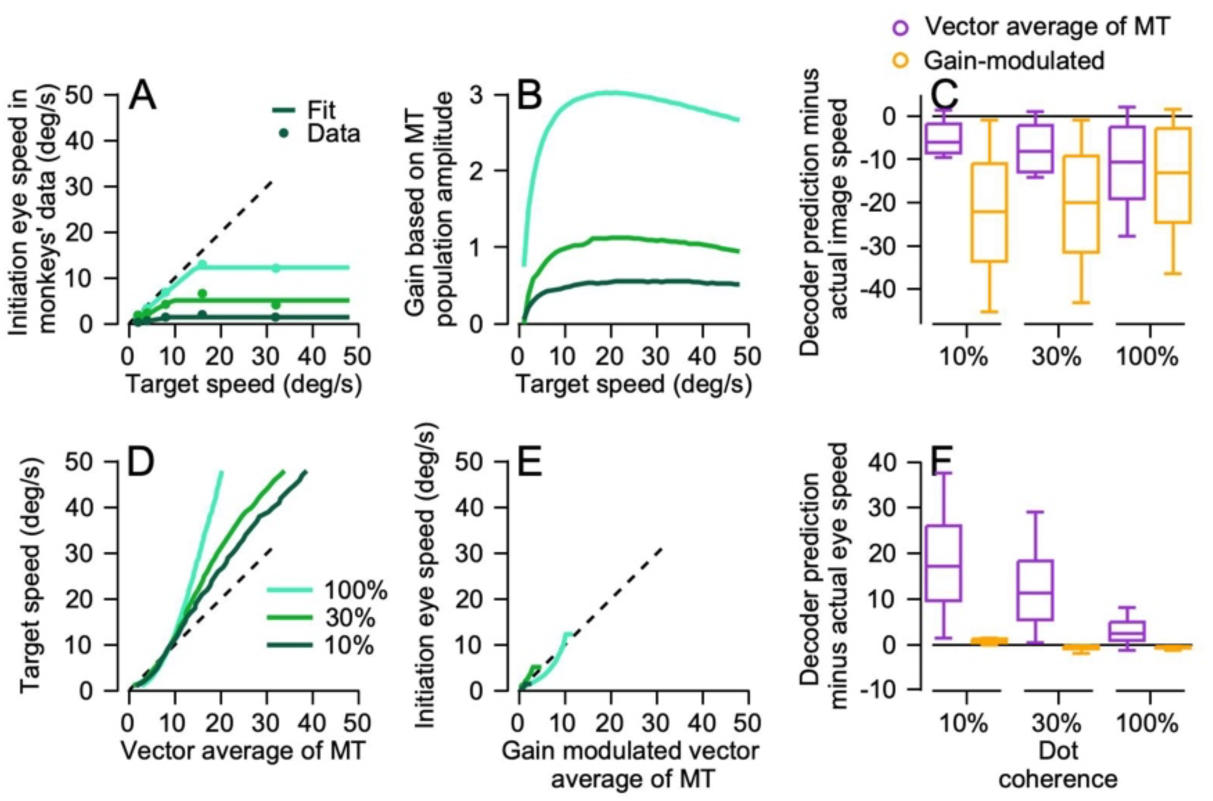
Decoding a simulated MT population response from the sum-of-Gaussians model to reproduce eye speed during pursuit initiation. **A**: Pursuit initiation eye speed in monkeys’ data as a function of the target for 3 dot coherences (100%, 30%, and 10%). Symbols plot the values from the data and the lines represent fits used to test predictions from decoding the sum-of-Gaussians population response. **B**: The gain of visual motor transformation based on the sum-of-Gaussians MT population amplitude. **C**: The difference of image speed and predictions of the two decoders. **D**: Target speed as a function of predictions from decoding by a vector average of the sum-of-Gaussians MT population response. **E**: Eye speed in monkeys’ data as a function of the gain-modulated vector average of the sum-of-Gaussians model of MT. **F:** The difference of eye speed and the predictions of the two decoders. In **A, B, D,** and **E**, cyan, green, and dark green lines show data and decoding performance for 100%, 30%, and 10% dot coherences. In **C** and **F**, purple and orange boxes and error bars show results for decoding based on an opponent vector average of MT versus a gain-modulated vector average.

Figures 8C and F summarize the predictions of the two decoders as a function of dot coherence and verify the impression from Figures 8D and E. The opponent vector average decoder (purple box and whiskers) predicts image speed fairly well and eye speed poorly, while the gain-modulated opponent vector average decoder (yellow box and whiskers) predicts eye speed well and image speed poorly. The results of decoding simulated MT responses support the accuracy of our model of pursuit. They also reinforce our conclusion that MT population responses capture the target speed reasonably well, but additional aspects of the sensory-motor decoder such as gain modulation from FEF_SEM_ are needed to account for the eye speeds in pursuit initiation.

### Effect of dot coherence and stimulus speed on the population response in MT and model population during steady-state pursuit

Inspection of Figure 2A and B reminds us of the conundrum during steady-state tracking: eye speed lags behind target speed, especially for lower dot coherences, so there is a non-zero image speed that fails to cause the eye to accelerate up to target speed. Now, we attempt to understand the basis for the conundrum by analyzing the activity of MT neurons during steady-state tracking.

A key difference exists between the visual conditions during pursuit initiation versus steady-state tracking because the eye moves along with the target during steady-state tracking. The difference between the target speed and eye speed produces an image speed on the fovea. For 100% coherence patches of dots, the difference between target and eye is minimal and the target is almost stationary relative to the fovea as the eye moves, but for lower dot coherences (or the highest target speeds at 100% coherence), eye speed is persistently slower than target speed and requires occasional saccades to keep the target foveated. Also, the background produces oppositely-directed motion across much of the retina. By analyzing MT activity during steady-state tracking, we sought to determine if the visual representation of motion emanating from MT during steady-state tracking represents image speed positively, in which case we would need to invoke additional downstream deficits that prevent the required eye acceleration to catch up to the target.

We characterized the relationship between image speed and change in eye speed during steadystate tracking by measuring firing rate in the interval from 600 to 660 ms after target motion onset and the change in eye speed in the interval from 660 to 780 ms after target motion onset. We used the change in eye speed as our metric for steady-state eye movement versus the absolute eye speed for the initiation of pursuit, but they are actually homologous because the eye speed during pursuit initiation is a change from zero deg/s. Change in eye speed is the correct metric because of considerable evidence that image motion causes eye acceleration (Lisberger and Westbrook 1985; Lisberger et al. 1981) and that zero image motion enables stable eye speeds (Morris and Lisberger 1987).

When the moving target was 100% coherence dots, targets speeds less than 16 deg/s caused almost perfect tracking so that image speed was small and change in eye speed negligible when target speed (Figure 9A, 4 of the 5 light green symbols). The one outlier symbol at an image speed of 10 deg/s and eye acceleration of –4 deg/s was for target speeds of 32 deg/s, when our monkeys did not sustain eye speed close to target speed even for 100% coherence dots. The data for low dot coherences agree qualitatively with the data for 100% coherence. In each case, especially the higher target speeds caused the eye to lag behind the target and we measured and modest negative changes in eye speed in spite of significant positive image speeds (Figure 9A). The relationship between change in eye speed and image speed was reasonably linear for each dot coherence, with decreasing slopes as dot coherence decreased, meaning higher image speeds at a given target speed (Figure 9A). We attribute the relatively poor tracking for target motion at 32 deg/s to the high density of slower target speeds in our stimulus set and its effect on the priors used by pursuit to control the gain of visual-motor transmission and therefore eye speed (Darlington et al. 2017).

**Figure 9:**
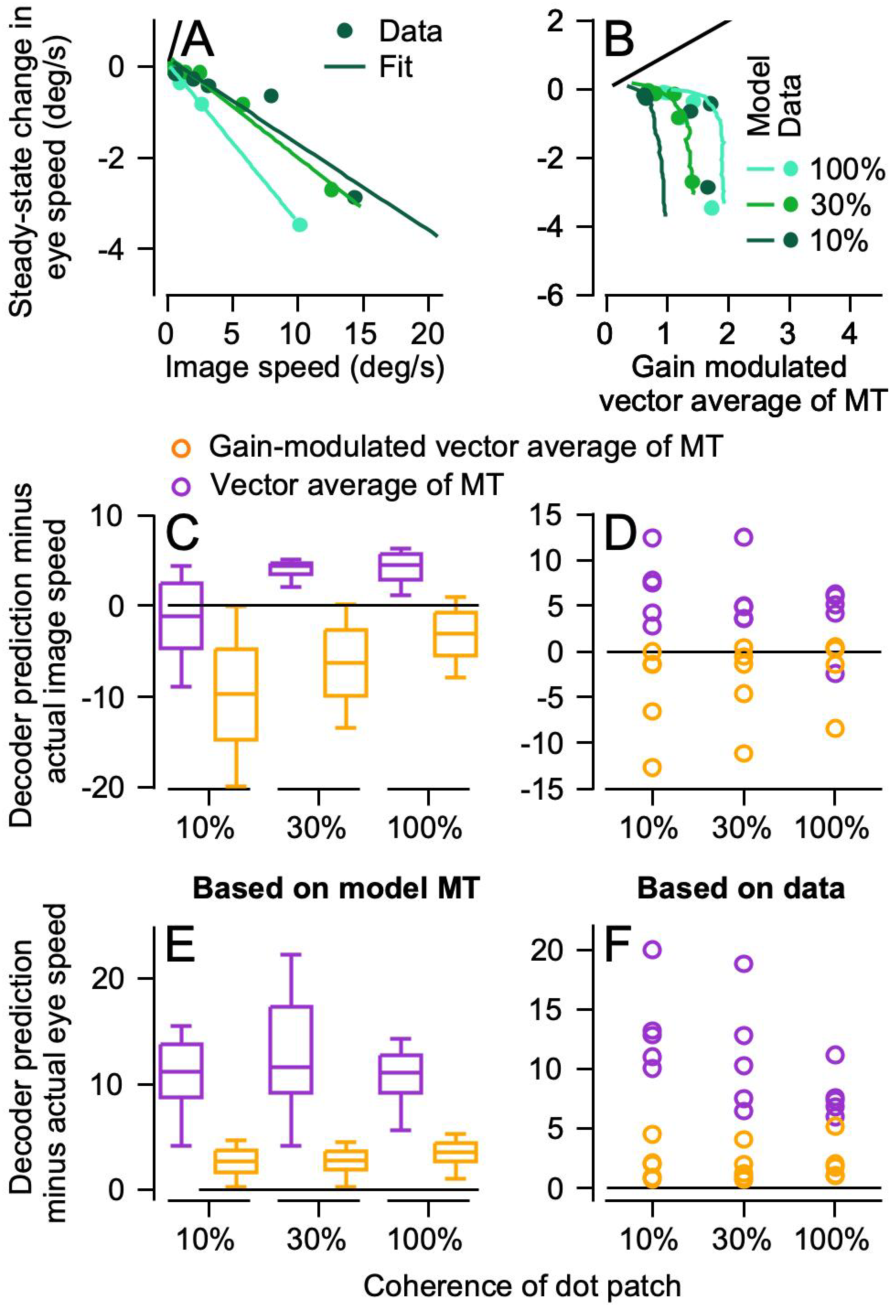
Decoding eye speed during steady-state tracking. **A**: Steady-state change in eye speed as a function of image speed. Symbols show the monkeys’ data and lines illustrate linear fits to the data. **B**: Steady-state change in eye speed versus the result of decoding with a gain-modulated vector average of MT. Symbols represent the values from decoding based on the data and traces show the results of decoding based on the extended sum-of-Gaussians MT population response. In **A** and **B**, cyan, green, and dark green show results for 100%, 30%, and 10% dot coherences. **C-F**: The difference between data and the predictions of decoders for based on the model population response (**C, E**) and the actual data (**D, F**). **C** and **D** show results for image speed; **E** and **F** show results for eye speed. In **C-F**, purple versus orange box and whiskers and symbols show results for decoding with opponent vector average of MT versus a gain-modulated vector average.

The same decoders used to predict image speed or (change in) eye speed successfully during pursuit initiation failed to capture either during steady-state tracking. The opponent vector average of MT activity during steady-state tracking overestimates image speed for most conditions (purple symbols, Figure 9C, D); it predicts large, positive eye speeds (purple symbols, Figure 9C, D) for both the data and model population responses. Gain-modulated vector averaging underestimates image speed severely (yellow symbols, Figure 9C, D) and predicts positive changes in eye speed when the eye is actually stable or decelerating (yellow symbols, Figure 9E, F). In summary, image speed during steady-state tracking is captured poorly by both decoders while even gain-modulated vector averaging consistently misestimates the sign of the change in eye speed, calling for eye acceleration when we observe eye deceleration.

We conclude that the visual signal is much more muddied during steady-state tracking when compared to the relative clarity during pursuit initiation. The disagreement between the predictions from decoding the MT population response and the actual eye movement data imply that MT is not the sole determinant of eye movement during steady-state tracking. Other mechanisms may be operating downstream and may be sensitive to the changes in motion reliability caused by reducing dot coherence. At the same time, the good agreement between the results of decoding the data versus the model MT population responses provides additional support for the accuracy of the sum-of-Gaussians modeling strategy.

### MT neurons respond appropriately to imposed image motion during steady-state tracking

One possible explanation for the pursuit system’s tolerance of persistent image motion during steady-state tracking is that MT is no longer responding to motion in a way that the pursuit system can use. Therefore, we next ask how individual neurons and the population in MT respond to perturbations that cause image speed during steady-state tracking and how those responses compare to what is expected given our data for pursuit initiation. We imposed pulses of target speed during steady-state tracking to afford explicit control of image speed.

Individual neurons in MT respond to pulses of target motion during steady-state tracking in a way that generally matched responses during pursuit initiation for similar image speeds. Figures 10A and B show the response of an example neuron to a target that initiated pursuit at 8 deg/s before increasing by 8 deg/s at 500 ms after motion onset. Comparison of the responses during pursuit initiation versus steady-state tracking reveals similar pulses of firing rate and similar changes in eye speed at each dot coherence, with larger responses for 100% vs. 10% coherence. In these recordings, we kept the overall image speed imposed during steady-state tracking the same across dot coherences by delivering smaller speed pulses at lower coherences. For example, we used speed pulses of 8 deg/s at 100% dot coherence, 5 deg/s for 30% dot coherence, and 3.5 deg/s for 10% dot coherence. Because of the persistent image motion during steady-state tracking of dots with lower coherences, systematic variation of the size of the speed pulse created nearly identical image speeds across dot coherence.

**Figure 10:**
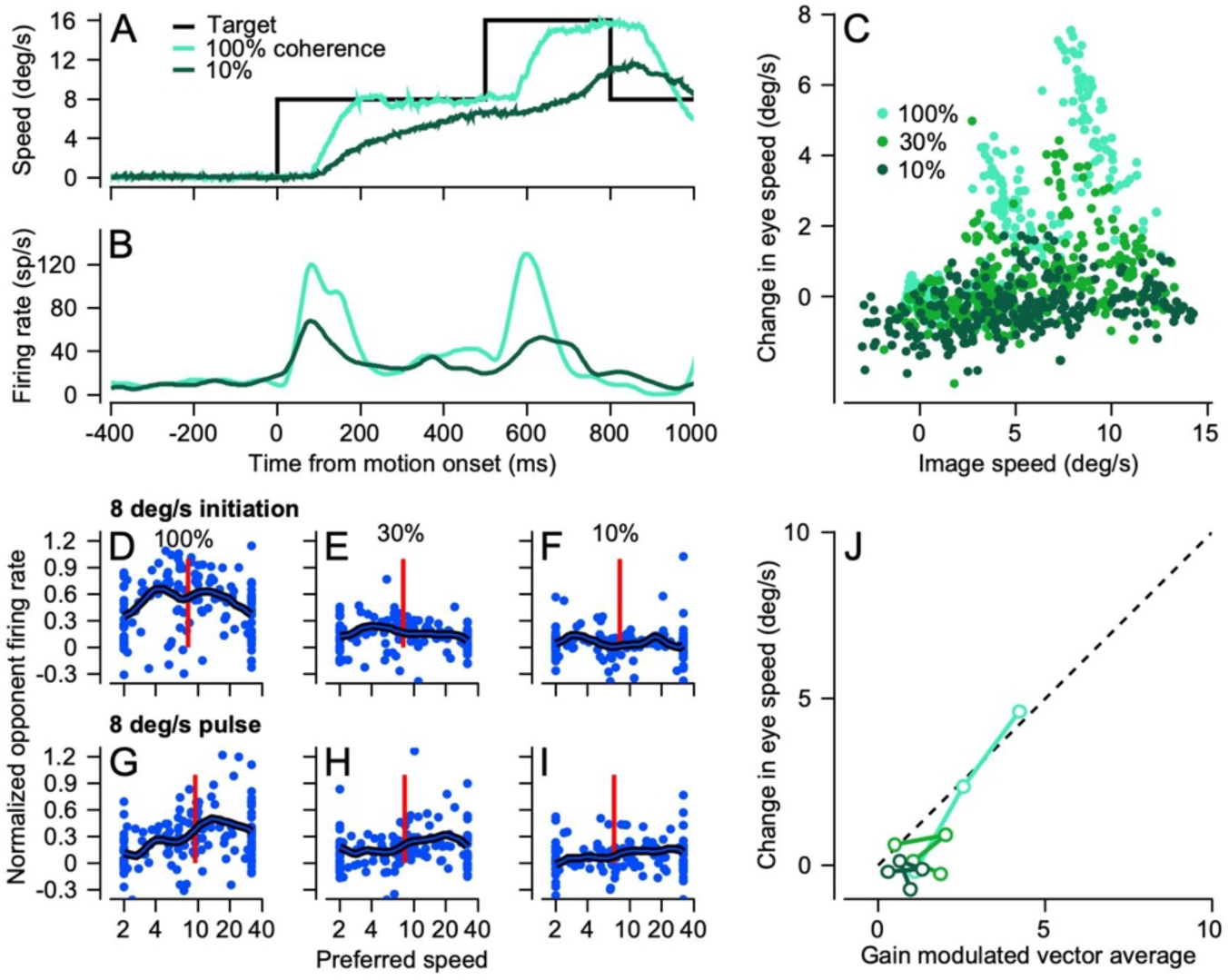
Responses of MT neurons to pulses of target speed during steady-state tracking. **A**: Example traces for eye and target speed. The black trace shows an 8 deg/s pulse of target speed from 500 to 800 ms after the onset of 8 deg/s target motion. **B**: Firing rate for an example neuron for the target and eye motions in **A**. In **A** and **B**, cyan and dark green traces show responses for 100% and 10% dot coherences. **C:** The change in eye speed caused by speed pulses as a function of the image speed. Each symbol shows the response in an average trial for a single experimental day during recordings from MT. Cyan, green, and dark green symbols show data for 100%, 30%, and 10% dot coherences. **D-I**: Normalized opponent population responses. Each symbol plots the response of one MT neuron as a function of its preferred speed. The thick trace shows the population average. The vertical red line indicates the image speed for each population response. The top and bottom rows of 3 graphs show data for pursuit initiation to a target of 8 deg/s and target pulses of 8 deg/s. From left to right, the three columns show data for dot coherences of 100%, 30%, and 10%. **J**: Comparison of change in eye speed from target speed pulses to the predictions from decoding the population response using the gain-modulated vector average decoder from Equation 5. Color scheme as in **C**.

The change in eye speed caused by speed pulses depended on both the image speed caused by the pulse and the coherence of the dot patch (Figure 10C). At any given image speed, responses were generally faster for higher dot coherences and in general the change in eye speed increased as a function of image speed. Here, we measured the change in eye speed from 60-120 ms after the onset of the speed pulse and the image speed (target minus eye speed) at the start of the pulse and plotted measurements from trial averages (Figure 10C). We conclude that eye acceleration is driven by visual motion, and presumably by MT, throughout pursuit.

Population responses in MT are similar (but not identical) for speed pulses during steady-state tracking and for pursuit initiation. For the motion of 100% coherence dots with an image speed of 8 deg/s, the population response for speed pulses (Figure 10G) is somewhat lower in amplitude than that for pursuit initiation (Figure 10D). Both population responses have a peak near 8 deg/s but the data for speed pulses show unexpectedly large responses in MT neurons with higher preferred speeds. Very similar trends appear in the data for 30% coherence dots (Figure 10E, H) and 10% coherence dots (Figure 10F, I). The eye speed predicted by the same gain-modulated vector average decoder that worked well for pursuit initiation recapitulates the eye speed caused by speed pulses fairly consistently across coherence and pulse speed (Figure 10J). We conclude that the population response in MT is still driving the pursuit system during steady-state tracking.

### MT shows machine-like responses to changes in dot coherence during steady-state tracking

Target motions that underwent increases or decreases in coherence during steady-state tracking revealed that eye speed and individual MT neurons respond with a machine-like quality. The approach of delivering changes in dot coherence during tracking allowed us to evaluate the effect of dot coherence alone, while prior image and eye speed are held constant. A change in coherence from 100% to 10% that occurs 500 ms after the onset of target motion (Figure 11A, black trace) causes eye speed to drop from that produced by continued motion 100% dot coherence (orange trace) to that produced by motion that started and continued with 10% dot coherence (green trace). At the same time, we observed parallel responses in the firing rate of a typical MT neuron (Figure 11B).

**Figure 11:**
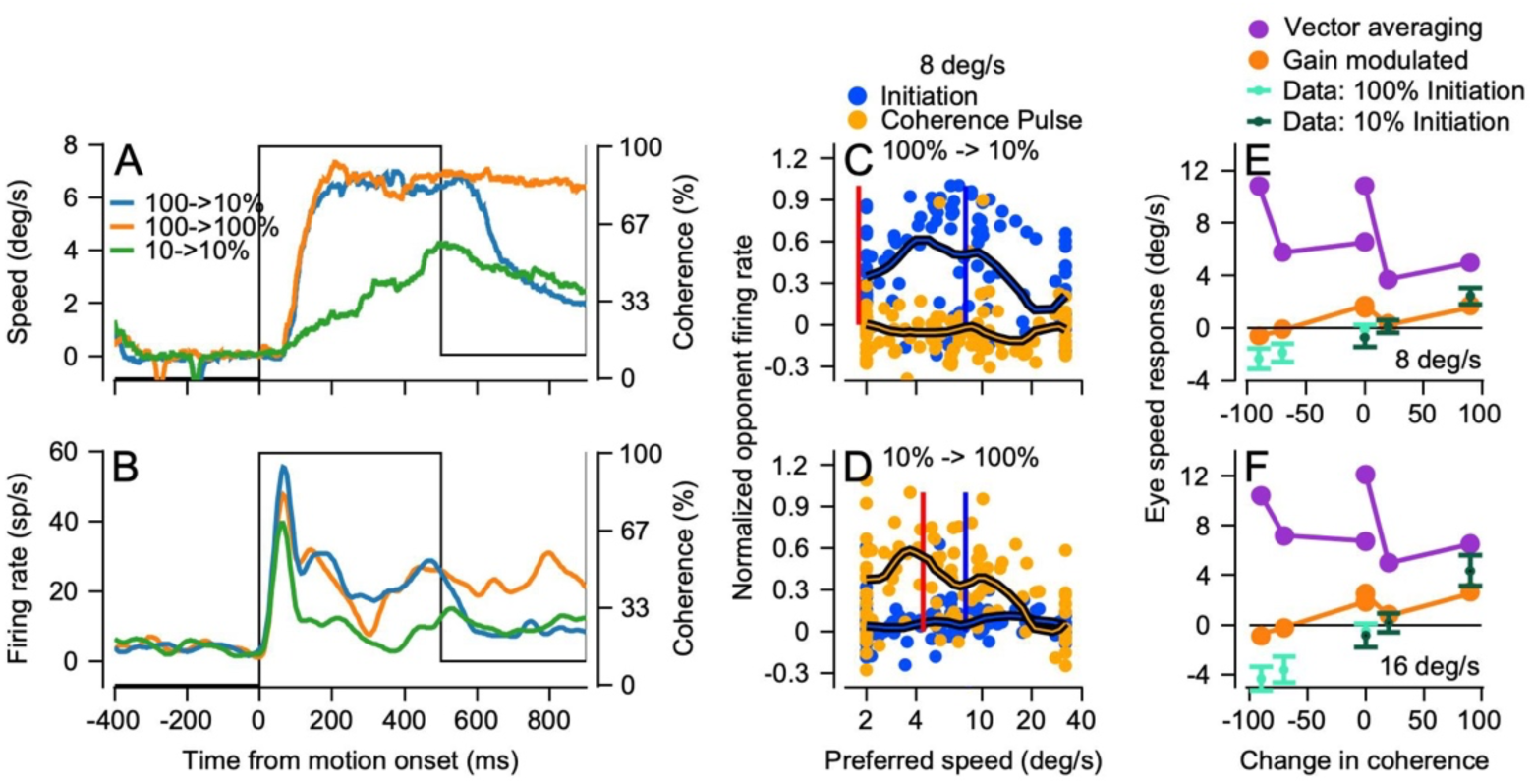
Effect of changes in dot coherence on the responses of MT neurons during steady-state tracking. **A**: Example eye speed traces as a function of time. Green and orange traces show responses to targets at 10% versus 100% dot coherence; blue trace shows responses when the coherence changes from 100% to 10% at 500 ms after the onset of target motion. **B**: Firing rate for an example neuron in response to the target motions in **A**. **C, D**: Population responses. Each symbol shows normalized opponent firing rate for different individual neurons in the population as a function of its preferred speed. Blue versus orange symbols show responses during pursuit initiation versus a change in coherence during steady-state tracking. The thick traces represent the population averages and the vertical blue and red lines represent the image speed during pursuit initiation and during the coherence pulse. **C:** Coherence drops from 100% to 10% at 8 deg/s. **D**: Coherence rises from 10% to 100% at 8 deg/s. **E, F**: Eye speed responses during steady-state tracking as a function of the change in coherence for targets at 8 deg/s (**E**) or 16 deg/s (**F**). The cyan and dark green small symbols show the eye speed response when the dot coherence started at 100% or 10%. Error bars represent standard deviation across experiments. Orange and purple symbols show the eye speed predicted from decoding the population response with gain-modulated vector averaging of MT versus with opponent vector averaging.

Population responses to increasing or decreasing dot coherence during steady-state tracking are also very machine-like. In the interval from 60-120 ms after decreases in dot coherence from 100 to 10%, the population responses were low in amplitude (Figure 11C), because coherence was low and the image motion that drove the response in the measurement interval was small (vertical red line). In contrast, the population responses were quite large after a change in dot coherence from 10 to 100% (Figure 11D), because coherence was high and image motion was greater than 4 deg/s. The population responses for changes in dot coherence (orange symbols) peak around the image speed present when the change in coherence occurs (vertical red line). Comparison of the population responses during pursuit initiation to those for coherence pulses is slightly tricky because of the differences in image speed, but in general we find reasonable agreement between the population responses for 100% dot coherence during pursuit initiation (Figure 11C, blue symbols) and for coherence steps from 10 to 100% (Figure 11D, orange symbols), with peaks at preferred speeds near image speed.

Decoding MT population activity and comparing to the eye acceleration shows that opponent vector averaging of the population response (Figure 11E, F, purple symbols) consistently overestimates the actual eye speed changes in the monkeys (small cyan and dark green symbols). Gain-modulated vector averaging of the population responses (Figure 11E, F, orange symbols) captures the change in eye speed for changes from low coherence to high coherence (positive values on x-axes), but consistently overestimates the change in eye speed for changes from 100% coherence to low coherence (negative values on x-axes). The latter finding is consistent with the results of decoding the population responses during steady-state tracking. When dot coherence is low, decoded MT population responses predict increases in eye speed, but the data show decreases in eye speed. We will argue below that another process is creating the eye deceleration.

### Why steady-state tracking asymptotes with eye speed stably much lower than target speed under some stimulus conditions

A model of pursuit that incorporates the effect of changing motion reliability through dot coherence helps us in two ways: 1) it tests our idea of how our observations in MT contribute to the visual drive for pursuit for high versus low dot coherence and 2) it formalizes our concepts of the balance that we predict between eye acceleration driven by the decoded visual drive from MT and eye deceleration resulting from deficits in motor pathways during low dot coherence. Thus, it explains how eye speed during steady-state tracking can be stable at speeds well below target speed in spite of active visual motion drive.

Our model (Figure 12, top diagram) is based on a control theory pursuit model proposed by Churchland et al. (2001) and expanded on by ourselves (Behling and Lisberger 2020). The model includes a feedforward path to simulate visual processing to create a command for eye acceleration that is modulated by a gain (g1). It uses a positive feedback loop of motor commands, with a gain (g2), to sustain eye speed in the absence of image motion, for example during perfect steady-state tracking. Dot coherence modulates the values of g1 and g2. We showed in our previous paper (Behling and Lisberger, 2020) that separate modulation of the two pathways can recreate the effects of dot coherence on the eye speeds observed during pursuit initiation and steady-state tracking.

**Figure 12:**
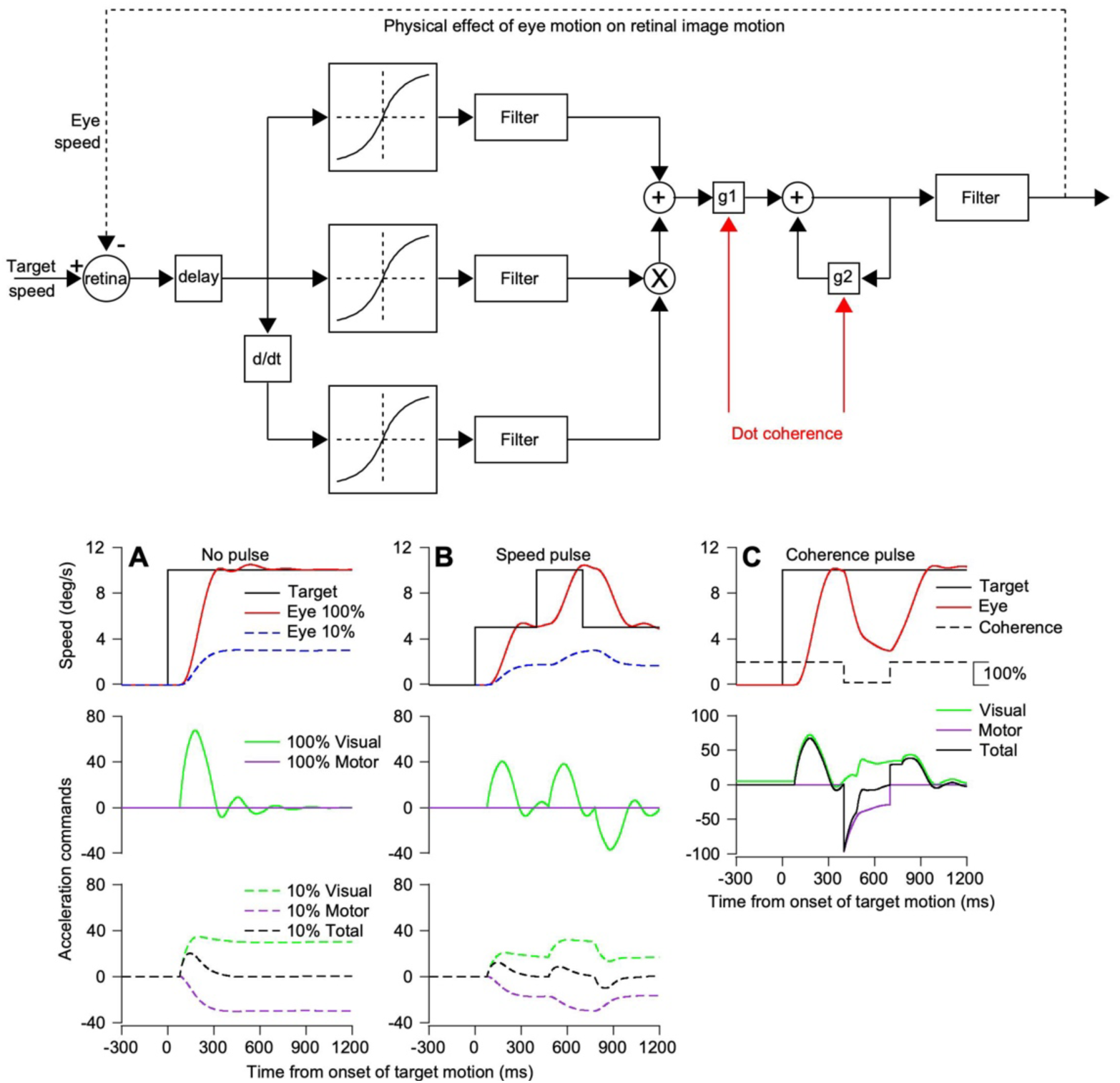
A control theory model of pursuit explains how steady-state eye speed can fail accelerate to target speed. The top diagram shows the control theory model used in Behling and Lisberger (2020) and derived from the model of Churchland and Lisberger (2001). **A-C**: The top row of graphs plots eye speed in degrees per second as a function of time from onset of target motion in milliseconds. Blue and red traces show eye speeds for targets with 100% and 10% coherence. The middle and bottom rows of graphs plot eye acceleration commands separately for visual and motor components of the model, as a function of time from onset of target motion. Black traces show target speed. Solid and dashed green traces show visual eye acceleration commands for 100% coherence and 10% coherence. Solid and dashed purple traces show the motor component of eye acceleration for 100% and 10% coherence. The dashed black trace shows the total acceleration command for 10% coherence. **A**: Initiation and steady-state pursuit without a pulse of speed or coherence. **B**: Response to a speed pulse of 5 deg/s during steady-state tracking. **C**: Response to a pulse of dot coherence (shown with the black dashed line in the upper panel).

For pursuit of a target that moves at constant speed, the output of the model recreated our behavioral observations with deficits due to dot coherence both at pursuit initiation and during steady-state tracking (Figure 12A). During pursuit of 100% coherence dots, the model’s eye speed accelerates to target speed and matches it through steady-state tracking. Eye speed is sustained perfectly by the positive feedback in the motor system, so that the visual drive for eye acceleration is transient (Figure 12A, continuous green trace) and the motor drive for eye acceleration or deceleration is nil. During pursuit of 10% dot coherence dots, both g1 and g2 are reduced so that the eye accelerates less and stabilizes at a speed considerably below target speed. Now, the visual drive for eye acceleration (dashed green trace) persists throughout steady-state tracking and the weakened positive feedback in the motor system creates a drive for eye deceleration (dashed purple trace) that is balanced perfectly by a visual drive that cannot accelerate the eye any further towards target speed.

The same phenomena play out when we deliver speed and coherence pulses, though transiently. The speed pulse (Figure 12B) drives eye acceleration comparable to that during pursuit. It is larger and more transient for 100% dot coherence than for 10% dot coherence. During steady-state pursuit, the visual drive perfectly balances the motor command for deceleration for 10% dot coherence. A pulse of target coherence from 100% to 10% and then back to 100% (Figure 12C, black dashed trace) creates a transient motor command for deceleration that dynamically balances with the subsequent visual drive for eye acceleration. Both return to normal when dot coherence returns to 100%.

The model explains why the eyes stabilize at speeds below target speed during pursuit of low coherence dots even though visual motion drive for eye acceleration is still present. This is reinforced by our recordings in MT that demonstrate speed representations appropriate to cause positive eye acceleration during steady-state tracking. Thus, the slow eye speeds during steady-state tracking are not from a misrepresentation of speed within the pursuit system but rather reflect the impact of poor motion reliability on both the sensory feedforward and motor feedback components of the system, in different ways that ultimately balance each other out.

## Discussion

We began this project with the goal of using changes in motion reliability to better understand how the representation of visual motion in area MT is transformed in downstream circuits to generate both the initiation and steady-state of smooth pursuit eye movements. In our previous paper (Behling and Lisberger 2020), we changed the coherence inside a patch of moving dots to control motion reliability and observed that reduced dot coherence causes persistent deficits in eye speed at the initiation of pursuit initiation as well as stable eye speeds that were slower than target speed during steady-state tracking. We were perplexed by the failure of the pursuit system to correct the persistent image motion during steady-state tracking. Thus, one of our major goals here was to determine why eye speeds slower than target speed were tolerated as well as why eye speed at the initiation of pursuit was affected by dot coherence. Our analysis provides an explanation for both effects, an explanation that is couched in terms of the functional organization of the sensory-motor decoder and its possible implementation in neural circuits.

Our strategy was to record from a major source of visual motion signals to drive pursuit, extrastriate area MT (Born et al. 2000; Groh et al. 1997; Newsome et al. 1985) and quantify how changes in dot coherence change the output of MT throughout the initiation and steady-state of a pursuit tracking movement. We then used our data to ask not how target motion is transformed into eye motion, but rather how the downstream sensory-motor decoder might decode output from MT into the observed initiation and steady-state of pursuit. Our approach changes the strategy for creating models of the pursuit system and thinking about how sensory signals are decoded by using the MT population response as the input to a sensory-motor decoder, instead of using the kinematics of target motion as the input.

We find that decrease dot coherence alters the amplitude but not the preferred speed of MT neurons during the initiation of pursuit. We also find that the responses of MT neurons during steady-state tracking are predicted poorly by their speed tuning during pursuit initiation, not a surprising finding given that MT neurons have complex responses to motion outside the classical receptive field (e.g. Born and Tootell, 1992) and the existence of large field image motion in the direction opposite to eye and target motion during steady-state tracking, because the eyes are moving across a stationary background. Because of the inability of speed tuning curves to predict MT responses during steady-state tracking, we developed a descriptive model of the full MT response during pursuit. Our “sum-of-Gaussians” model is not intended to reproduce any mechanisms of MT responses, but rather to provide an accurate description that can be used in future computational models of the pursuit circuit. The fact that we obtained essentially the same answers from decoding the population response from actual data and a larger, more homogeneous model population response implies that our model provides a reasonable description of the actual MT population response.

Our laboratory has developed a concept of the sensory-motor decoder for pursuit that is based on the anatomy of the pursuit system and evidence that parallel downstream pathways perform different computations based on the sensory representation of motion in MT (Egger and Lisberger 2022). Armed with quantitative measurements of the MT population response and the evoked eye movements for a range of target speeds and motion reliabilities (i.e. dot coherences), we now can make a quantitative statement about how the sensory-motor decoder works. Importantly, our conclusions about the sensory-motor decoder that are better aligned with the biology of the system compared to attempts to characterize decoders in terms of equations such as vector averaging (Churchland and Lisberger 2001b; Groh et al. 1997) or concepts like Bayesian inference (Darlington et al. 2017), Kalman filters (Orban de Xivry et al. 2013), or probabilistic population codes (Ma et al. 2006).

Figure 13 summarizes our conclusions about the sensory-motor decoder for pursuit. To account for the transformation between the MT population responses we measured and the evoked pursuit eye movements, we required three pathways and two sites of modulation by motion reliability. One pathway, shown in blue, finds the preferred speed at the peak of the MT population response and estimates the image speed the system is seeing. A second pathway, shown in green, measures the amplitude of the MT population response and uses it to create a representation of sensorymotor gain in the smooth eye movement region of the frontal eye fields (FEF or FEF_SEM_). Downstream, presumably in the pons (Darlington and Lisberger 2022), the output from FEF_SEM_ modulates the speed estimate from MT to create eye speed at pursuit initiation that increases as a function of target speed and decreases as a function of dot coherence. The combination of the first and second pathways in our conceptual framework explains why reduced dot coherence does not shift the peak of the MT population response but reduces its amplitude and thereby causes decreased eye speed in the initiation of pursuit. It also would account for the fact that low contrast targets move the peak of the MT population response towards faster preferred speeds while reducing the amplitude of the population response (Krekelberg et al. 2006) and therefore cause reduced eye speed in the initiation of pursuit (Darlington et al. 2017; Lisberger and Westbrook 1985).

**Figure 13:**
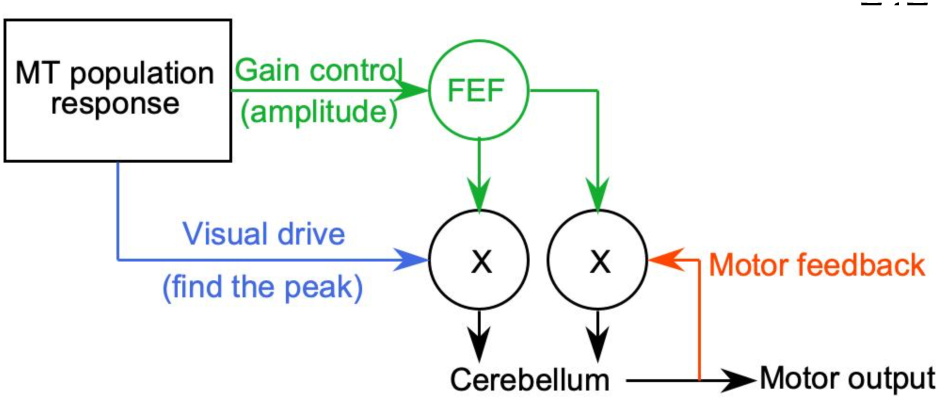
Conceptual model of the sensory-motor decoder for pursuit. Green pathways show the control of sensory-motor gain, blue pathway shows visual drive by estimates of target speed, red pathway shows motor feedback that sustains steady-state eye speed. The circles with “X” inside represent sites of multiplication where gain modulates other signals.

We required a third pathway to account for pursuit eye speeds during steady-state tracking, especially to account for the fact that eye speeds settled well below target speed for low values of dot coherence. Because the output of MT was appropriate to command positive eye acceleration during steady-state tracking, decoders that successfully predicted eye speed during pursuit initiation failed during steady-state tracking. They predicted eye acceleration that did not appear in the data. Despite the persistent and often large image speed during steady-state tracking, the eyes either remain stable or decelerate. We demonstrated that MT was still involved in driving pursuit during steady-state tracking by using speed pulses to verify the responsiveness of MT to image speed during steady-state tracking. We also used changes in dot coherence pulses to verify that MT had machine-like responses to dot coherence during steady-state tracking. We conclude that area MT continues to provide viable visual motion drive during steady-state tracking that should be driving the eye speed up to target speed. Thus, a third pathway was needed.

In Figure 13, the third pathway uses extraretinal signals related to eye movement as positive feedback to sustain eye speed. Corollary discharge through the floccular complex of the cerebellum has long been thought to be the substrate of the positive feedback (Lisberger and Fuchs 1978a; Stone and Lisberger 1990). To account for the paradox that the output from MT during steady-state tracking of low-coherence dots should, but does not, drive eye acceleration up to target speed, we propose that the visual-motor gain signal from FEF_SEM_ also controls the gain of the positive feedback. Then, reduced dot coherence should cause a tendency for steady-state eye speed to decelerate, meaning that the eye acceleration command from MT needs to counteract the deceleration merely to keep eye speed steady at less than target speed. Our control theory model shows the plausibility of this explanation, even though biological details will need to be the topic for future research.

The most parsimonious explanation for the deficits in pursuit during steady-state tracking of low dot coherence targets is that the visual drive signal still is present and used by the pursuit system and the downstream motor system is modulated instead. We anticipate that the same explanation might account for deficits in steady-state tracking following lesions of the medial superior temporal area, MST, (Dürsteler and Wurtz 1988), FEF_SEM_ (Shi et al. 1998), and the floccular complex of the cerebellum (Rambold et al. 2002), or even for the smooth eye movement deficit in schizophrenia (MacAvoy and Bruce 1995). Thus, our analysis explains the tolerance of the pursuit system for persistent image motion during steady-state tracking based on visual-motor gain that decreases visual drive paired with loss of commands that normally sustain eye speed automatically. We propose that there is parallel modulation of the sensory and motor paths before they integrate to drive eye movements. Our findings build on previous work and provide a novel perspective in the nuances and subtleties of sensory-motor control in an accessible and well-known system. Future investigation of the responses in FEF_SEM_ and the floccular complex of the cerebellum will help pin down the implementation of our observed parallel modulation.

## Acknowledgements

We thank Stefanie Tokiyama and Bonnie Bowell for technical assistance, and J. Patrick Mayo for guidance in mapping MT. Research supported by NIH grant EY027373.

## References

1. Behling S, Lisberger SG. Different mechanisms for modulation of the initiation and steady-state of smooth pursuit eye movements. J Neurophysiol 123: 1265–1276, 2020.

2. Born RT, Groh JM, Zhao R, Lukasewycz SJ. Segregation of object and background motion in visual area MT: effects of microstimulation on eye movements. Neuron 26: 725–734, 2000.

3. Born RT, Tootell RB. Segregation of global and local motion processing in primate middle temporal visual area. Nature 357: 497–499, 1992.

4. Britten KH, Shadlen MN, Newsome WT, Movshon JA. The analysis of visual motion: a comparison of neuronal and psychophysical performance. J Neurosci 12: 4745–4765, 1992.

5. Britten KH, Shadlen MN, Newsome WT, Movshon JA. Responses of neurons in macaque MT to stochastic motion signals. Vis Neurosci 10: 1157–1169, 1993.

6. Churchland MM, Lisberger SG. Apparent motion produces multiple deficits in visually guided smooth pursuit eye movements of monkeys. J Neurophysiol 84: 216–235, 2000.

7. Churchland MM, Lisberger SG. Shifts in the population response in the middle temporal visual area parallel perceptual and motor illusions produced by apparent motion. J Neurosci 21: 9387–9402, 2001a.

8. Churchland MM, Lisberger SG. Experimental and computational analysis of monkey smooth pursuit eye movements. J Neurophysiol 86: 741–759, 2001b.

9. Darlington TR, Beck JM, Lisberger SG. Neural implementation of Bayesian inference in a sensorimotor behavior. Nat Neurosci 21: 1442–1451, 2018.

10. Darlington TR, Lisberger SG. Sensory-motor computations in a cortico-pontine pathway. BioRxiv, 2022. doi:10.1101/2022.02.22.481514.

11. Darlington TR, Tokiyama S, Lisberger SG. Control of the strength of visual-motor transmission as the mechanism of rapid adaptation of priors for Bayesian inference in smooth pursuit eye movements. J Neurophysiol 118: 1173–1189, 2017.

12. Desimone R, Ungerleider LG. Multiple visual areas in the caudal superior temporal sulcus of the macaque. J Comp Neurol 248: 164–189, 1986.

13. Dubner R, Zeki SM. Response properties and receptive fields of cells in an anatomically defined region of the superior temporal sulcus in the monkey. Brain Res 35: 528–532, 1971.

14. Dürsteler MR, Wurtz RH. Pursuit and optokinetic deficits following chemical lesions of cortical areas MT and MST. J Neurophysiol 60: 940–965, 1988.

15. Egger SW, Lisberger SG. Neural structure of a sensory decoder for motor control. Nat Commun 13: 1829, 2022.

16. Groh JM, Born RT, Newsome WT. How is a sensory map read Out? Effects of microstimulation in visual area MT on saccades and smooth pursuit eye movements. J Neurosci 17: 4312–4330, 1997.

17. Hall NJ, Herzfeld DJ, Lisberger SG. Evaluation and resolution of many challenges of neural spike sorting: a new sorter. J Neurophysiol 126: 2065–2090, 2021.

18. Krekelberg B, van Wezel RJA, Albright TD. Interactions between speed and contrast tuning in the middle temporal area: implications for the neural code for speed. J Neurosci 26: 8988–8998, 2006.

19. Lisberger SG, Evinger C, Johanson GW, Fuchs AF. Relationship between eye acceleration and retinal image velocity during foveal smooth pursuit in man and monkey. J Neurophysiol 46: 229–249, 1981.

20. Lisberger SG, Fuchs AF. Role of primate flocculus during rapid behavioral modification of vestibuloocular reflex. I. Purkinje cell activity during visually guided horizontal smooth-pursuit eye movements and passive head rotation. J Neurophysiol 41: 733–763, 1978a.

21. Lisberger SG, Fuchs AF. Role of primate flocculus during rapid behavioral modification of vestibuloocular reflex. II. Mossy fiber firing patterns during horizontal head rotation and eye movement. J Neurophysiol 41: 764–777, 1978b.

22. Lisberger SG, Movshon JA. Visual motion analysis for pursuit eye movements in area MT of macaque monkeys. J Neurosci 19: 2224–2246, 1999.

23. Lisberger SG, Westbrook LE. Properties of visual inputs that initiate horizontal smooth pursuit eye movements in monkeys. J Neurosci 5: 1662–1673, 1985.

24. Lisberger SG. Visual guidance of smooth-pursuit eye movements: sensation, action, and what happens in between. Neuron 66: 477–491, 2010.

25. Lisberger SG. Visual guidance of smooth pursuit eye movements. Annu Rev Vis Sci 1: 447–468, 2015.

26. MacAvoy MG, Bruce CJ. Comparison of the smooth eye tracking disorder of schizophrenics with that of nonhuman primates with specific brain lesions. Int J Neurosci 80: 117–151, 1995.

27. Maunsell JH, Van Essen DC. Functional properties of neurons in middle temporal visual area of the macaque monkey. I. Selectivity for stimulus direction, speed, and orientation. J Neurophysiol 49: 1127–1147, 1983.

28. Ma WJ, Beck JM, Latham PE, Pouget A. Bayesian inference with probabilistic population codes. Nat Neurosci 9: 1432–1438, 2006.

29. Miles FA, Fuller JH. Visual tracking and the primate flocculus. Science 189: 1000–1002, 1975.

30. Morris EJ, Lisberger SG. Different responses to small visual errors during initiation and maintenance of smooth-pursuit eye movements in monkeys. J Neurophysiol 58: 1351–1369, 1987.

31. Newsome WT, Wurtz RH, Dürsteler MR, Mikami A. Deficits in visual motion processing following ibotenic acid lesions of the middle temporal visual area of the macaque monkey. J Neurosci 5: 825–840, 1985.

32. Orban de Xivry J-J, Coppe S, Blohm G, Lefèvre P. Kalman filtering naturally accounts for visually guided and predictive smooth pursuit dynamics. J Neurosci 33: 17301–17313, 2013.

33. Osborne LC, Lisberger SG. Spatial and temporal integration of visual motion signals for smooth pursuit eye movements in monkeys. J Neurophysiol 102: 2013–2025, 2009.

34. Pack CC, Hunter JN, Born RT. Contrast dependence of suppressive influences in cortical area MT of alert macaque. J Neurophysiol 93: 1809–1815, 2005.

35. Priebe NJ, Cassanello CR, Lisberger SG. The neural representation of speed in macaque area MT/V5. J Neurosci 23: 5650–5661, 2003.

36. Priebe NJ, Lisberger SG. Estimating target speed from the population response in visual area MT. J Neurosci 24: 1907–1916, 2004.

37. Rambold H, Churchland A, Selig Y, Jasmin L, Lisberger SG. Partial ablations of the flocculus and ventral paraflocculus in monkeys cause linked deficits in smooth pursuit eye movements and adaptive modification of the VOR. J Neurophysiol 87: 912–924, 2002.

38. Ringach DL. A “tachometer” feedback model of smooth pursuit eye movements. Biol Cybern 73: 561–568, 1995.

39. Robinson DA, Gordon JL, Gordon SE. A model of the smooth pursuit eye movement system. Biol Cybern 55: 43–57, 1986.

40. Robinson DA. The mechanics of human smooth pursuit eye movement. J Physiol (Lond) 180: 569–591, 1965.

41. Shi D, Friedman HR, Bruce CJ. Deficits in smooth-pursuit eye movements after muscimol inactivation within the primate’s frontal eye field. J Neurophysiol 80: 458–464, 1998.

42. Stanton GB, Friedman HR, Dias EC, Bruce CJ. Cortical afferents to the smooth-pursuit region of the macaque monkey’s frontal eye field. Exp Brain Res 165: 179–192, 2005.

43. Stone LS, Lisberger SG. Visual responses of Purkinje cells in the cerebellar flocculus during smooth-pursuit eye movements in monkeys. I. Simple spikes. J Neurophysiol 63: 1241–1261, 1990.

44. Tanaka M, Lisberger SG. Role of arcuate frontal cortex of monkeys in smooth pursuit eye movements. I. Basic response properties to retinal image motion and position. J Neurophysiol 87: 2684–2699, 2002a.

45. Tanaka M, Lisberger SG. Enhancement of multiple components of pursuit eye movement by microstimulation in the arcuate frontal pursuit area in monkeys. J Neurophysiol 87: 802–818, 2002b.

46. Zimmerman DW. Invalidation of parametric and nonparametric statistical tests by concurrent violation of two assumptions. The Journal of Experimental Education 67: 55–68, 1998.

